# Ensemble Switching Unveils a Kinetic Rheostat Mechanism of the Eukaryotic Thiamine Pyrophosphate Riboswitch

**DOI:** 10.1101/2021.03.12.434875

**Authors:** Junyan Ma, Nabanita Saikia, Subash Godar, George L. Hamilton, Feng Ding, Joshua Alper, Hugo Sanabria

## Abstract

Thiamine pyrophosphate (TPP) riboswitches regulate thiamine metabolism by inhibiting the translation of enzymes essential to thiamine synthesis pathways upon binding to thiamine pyrophosphate in cells across all domains of life. Recent work on the *Arabidopsis thaliana* TPP riboswitch suggests a multi-step TPP binding process involving multiple riboswitch conformational ensembles and that Mg^2+^ dependence underlies the mechanism of TPP recognition and subsequent transition to the translation-inhibiting state of the switching sequence followed by changes in the expression platform. However, details of the relationship between TPP riboswitch conformational changes and interactions with TPP and Mg^2+^ in the aptamer domain constituting this mechanism are unknown. Therefore, we integrated single-molecule multiparameter fluorescence and force spectroscopy with atomistic molecular dynamics simulations and found that conformational transitions within the aptamer domain associated with TPP and Mg^2+^ ligand binding occurred between at least five different ensembles on timescales ranging from μs to ms. These dynamics are at least an order of magnitude faster than folding and unfolding kinetics associated with translation-state switching in the switching sequence. Moreover, we propose that two pathways exist for ligand recognition. Together, our results suggest a dynamic ensemble switching of the aptamer domain that may lead to the translation-inhibiting state of the riboswitch. Additionally, our results suggest that multiple configurations could enable inhibitory tuning manifested through ligand-dependent changes via ensemble switching and kinetic rheostat-like behavior of the *Arabidopsis thaliana* TPP riboswitch.

## INTRODUCTION

Riboswitches, first reported by Henkin (1) and later named by Breaker and co-workers (2), are mRNA elements that bind small metabolites to regulate transcription or translation machinery in a cis-fashion (3–8). Typically, two distinct functional domains, including the ligand-sensing aptamer domain and the transcription or translation regulating expression platform, compose riboswitches. The aptamer domain recognizes an effector molecule, or ligand, causing structural changes in the expression platform domain that ultimately regulates gene expression. Aptamer domains can adopt multiple three-dimensional configurations, allowing for target recognition and further trapping of the ligand in the bound state (9). Therefore, riboswitches require high selectivity in target ligand recognition to elicit the appropriate regulatory response.

The thiamine pyrophosphate (TPP) riboswitch, also known as Thi-box riboswitch, regulates the transport and synthesis of thiamine and its phosphorylated derivatives (10) in response to changes in TPP concentration. TPP is the active form of vitamin B_1_, which is an essential coenzyme in many archaea, bacteria, and eukaryotes (11–14). Moreover, the TPP riboswitch has been postulated as an important therapeutic target for novel antibacterial drugs (15). For example, pyrithiamine pyrophosphate (PTPP, an antimicrobial agent) downregulates essential thiamine production in *Bacillus subtilis* and *Aspergillus oryzae*, demonstrating the potential use of riboswitches as targets for antibacterial or antifungal drug design (16).

The TPP riboswitch consists of five base-paired helices (Figure 1A, P1 to P5 (17)). The TPP riboswitch’s aptamer domain comprises two sensor helix arms, each composed of two stacked helices connected through a bulge. The stacking of helices P2 and P3 forms the pyrimidine sensor helix, and the stacking of helices P4 and P5 forms the pyrophosphate sensor helix domain (18). The TPP riboswitch’s expression platform comprises a secondary structural switch that interfaces with the translational machinery through species dependent mechanisms (e.g., prevents or allows ribosome binding to the expression platform). This expression platform is connected to the aptamer domain at the 3’ end of the P1 switch helix at the junction of the sensor arms (Figure 1A). The long-standing model of the TPP riboswitch’s mechanism involves two mutually exclusive structural conformations in the aptamer domain (Figure 1B) corresponding to the “ON” and “OFF” states of the riboswitch’s expression platform that enable and inhibit translation of enzymes essential to thiamine synthesis pathways, respectively. The model suggests that the expression platform’s OFF state corresponds to the aptamer domain’s TPP-bound state, while the ON state corresponds to the ligand-free state (5). Further, the ON/OFF behavior in similar riboswitches, such as *E. coli* ThiM (19,20) is coupled to formation of the switch helix P1, with the formed helix corresponding to the OFF state and unpairing of P1 corresponding to the ON state. This is because the 3’ bases of the P1 helix comprise the anti-anti-Shine-Dalargo (SD) sequence for the anti-SD and SD sequences found in the expression platform. Unpairing of P1 allows sequestering of the anti-SD by the anti-anti-SD in P1 and leaves the SD riboswitch binding site available for ribosome interactions. Hence, we refer to the anti-anti-SD sequence of P1 as the switching sequence. However, implicit RNA dynamics suggest the possibility of an ensemble of configurations rather than a small number of discrete states. Both binary ON/OFF and ensemble-like tuning behavior have been observed for various riboswitches, in some cases with differing behavior for the same riboswitch in different organisms (21).

**Figure 1.**
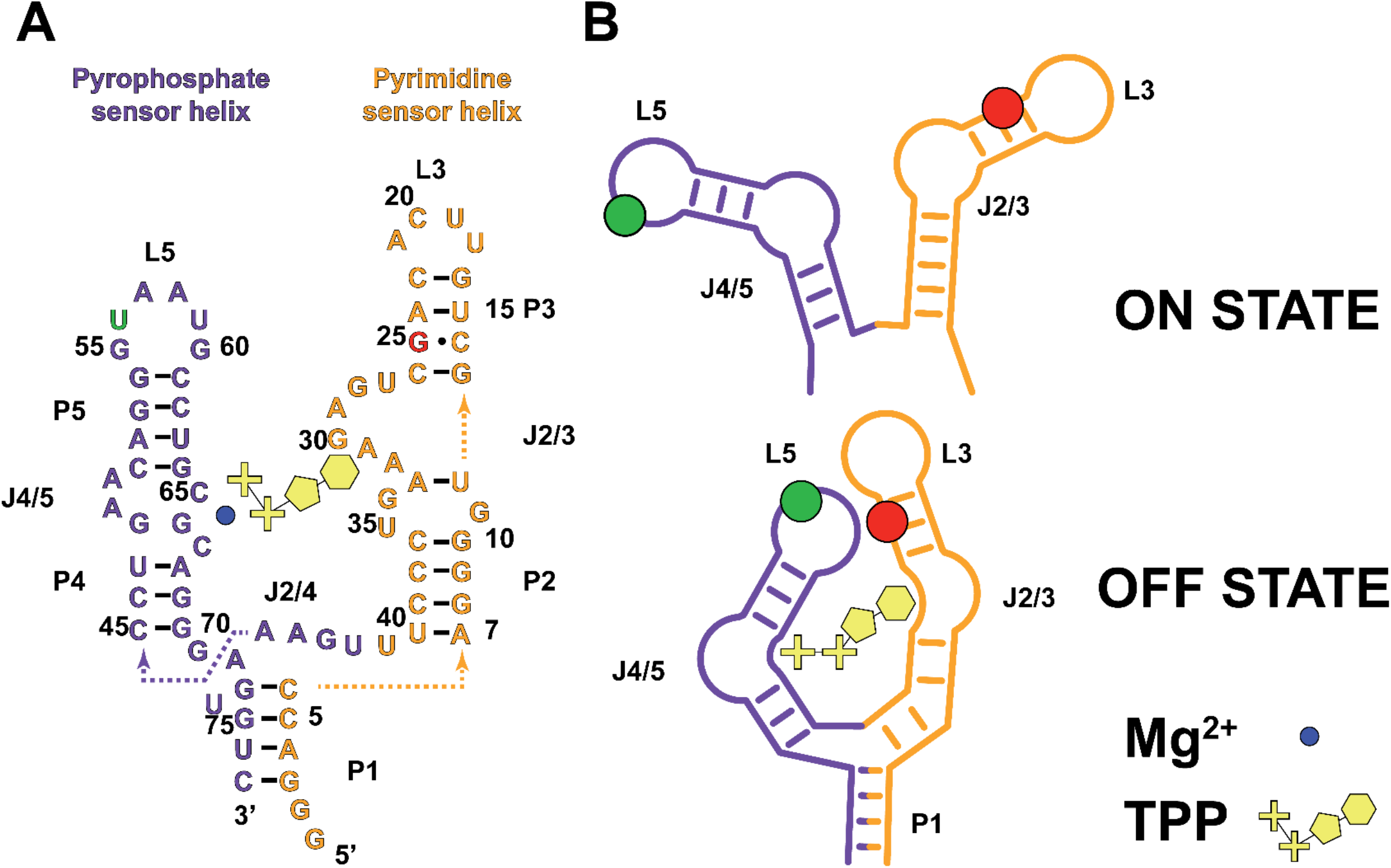
Model of TPP binding to the aptamer domain of the Arabidopsis thaliana TPP riboswitch. A) Primary and secondary structure schematic of the aptamer domain of TPP riboswitch used in all experiments and simulations, referred as TPP riboswitch throughout. Five Watson-Crick base-paired (black) helices (P1-P5) connect through three bulges (J2/3, J2/4, and J4/5) and two loops (L3 and L5). The two helices of the aptamer domain (P4/P5, purple and P2/P3, orange) sense the pyrophosphate (pluses) and pyrimidine (ring structure) ends of the TPP ligand (yellow), respectively, and a coordinating Mg^2+^ ion (blue circle). The P1 helix prior to the expression platform emerges from J2/4 to form a three-way junction and comprises the switching sequence that controls the “ON” or “OFF” behavior of the riboswitch. We truncated the sequence of L3 to match the crystal structure and added donor (Alexa 488 in position 56, green) and acceptor (Cy-5 in position 25, red) fluorophores (See Methods for details on riboswitch design, (17)) to L5 and near L3, respectively. B) Schematic of the TPP riboswitch “ON” and “OFF” states. Current models of the TPP riboswitch mechanism suggest that the TPP riboswitch is in the “ON state” (top) when the fluorescent markers (green circle for the donor, red circle for the acceptor, see Methods) are far apart (low FRET) and that the TPP riboswitch is in the “OFF state” (bottom) when the fluorescent markers are close together (high FRET) upon TPP binding between J4/5 and J2/3 in the aptamer domain. Figure modified from (17).

Previous work on TPP riboswitches has focused on the TPP ligand recognition process. Typically, TPP binds to TPP riboswitches with high affinity (up to 20 nM) (5,22) because the stacking of the phosphate sensor helix over the pyrimidine sensor helix allows recognition of the pyrimidine ring and pyrophosphate group of TPP (17,23,24). However, the situation is complicated because TPP binding is coordinated by a Mg^2+^ ion (25,26). Further complication arises because dynamic equilibrium exists between multiple aptamer domain conformational states, even when bound to TPP (18,21,26). Additionally, P1 helix formation of the switching sequence at the junction with the expression platform is necessary to further stabilize TPP binding to the aptamer domain and to transition to the OFF state (19). Moreover, recent experiments suggest a two-step TPP binding process, going from a weakly-bound intermediate configuration to the tightly-bound final state (27,28).

In the TPP riboswitch, it is thought that the two sensor helices close upon recognition of the pyrimidine and pyrophosphate groups of TPP (17). However, the configuration of the aptamer domain upon target recognition, but before it fully closes, remains unclear. It has been suggested that J4/5 kinks to facilitate the Mg^2+^-coordinated binding of the TPP’s phosphates, while the J2/3 bulge quasi-simultaneously kinks to accommodate the pyrimidine base of TPP on the other domain (17). However, this mechanism of binding remains somewhat speculative because these conformational transitions are thought to be rather subtle and fast (sub-millisecond); thus, they are challenging to characterize. Nonetheless, there are some reports of slow (second timescales) dynamic transitions, which would require that the transitions have high energetic cost (19). Similar riboswitches exhibit slow dynamics, such as the co-transcriptional folding and dynamics of the *E. coli* thiM riboswitch, which has 68% identity to the *Arabidopsis thaliana* TPP-riboswitch, that were revealed by single-molecule FRET (smFRET) (29).

To solve the apparent discrepancy between fast and slow mechanisms as well as to probe the ON and OFF configurations and potential intermediate states, we integrated single-molecule Multiparameter Fluorescence Spectroscopy (smMFS) (30), including time-correlated single-photon counting (TCSPC) (31), calculation of fluorescence parameters for single-molecule events (burst-integrated) (32), and fluorescence fluctuation methods using a the FRET labeled aptamer domain of the *Arabidopsis thaliana* TPP riboswitch (33,34), with all-atom discrete molecular dynamic (DMD) simulations (35) and optical tweezer force spectroscopy measurements. We found that the *Arabidopsis thaliana* TPP riboswitch’s aptamer domain exhibits Mg^2+^- and TPP-dependent ensemble switching that give rise to two distinct binding pathways in the transition to the known OFF state with both Mg^2+^ and TPP bound. Further, we only observe the TPP bound configuration similar to the crystallographic structure at extreme conditions of saturating Mg^2+^ and TPP, while in moderate and physiological conditions transitions OFF-like states and other intermediates are transiently populated at sub millisecond timescales. Together, our results suggest the TPP riboswitch’s aptamer domain is a conformational switching capable of tuning function through ligand-dependent dynamics as a kinetic rheostat rather than a binary all or nothing switch.

## MATERIAL AND METHODS

### TPP riboswitch design

The sequence of *Arabidopsis thaliana* TPP riboswitch aptamer domain was truncated to match the crystal structure (17) and to directly compare our results with prior structural information. To determine the best locations for fluorophore labeling and to maximize the resolution of expected conformational changes, we ran a coarse-grained simulation (36) whose trajectories were post-processed for modeling of the fluorescent markers at all possible pair distances. We calculated inter-label distances at each point in the trajectory by sampling accessible volumes generated for coarse-grained dyes linked in silico to the RNA backbone, as previously described (37). Based on all possible inter-label distances, we selected positions 25 and 56 for maximum resolvability of distance changes between two identified configurations from the coarse-grained simulations (Supplemental Information (SI) Figure S1). Using orthogonal click chemistry, the two selected nucleotides (Y(25)= 2′-dG-(N2-C6-aminoCy5) and X(56)= rU(2′-O-propargyl Al 488)) were incorporated into the synthesized TPP riboswitch sequence, 5’-GGG ACC AGG GGU GCU UGU UCA CAG **Y**CU GAG AAA GUC CCU UUG AAC CUG AAC AGG G**X**A AUG CCU GCG CAG GGA GUG UC-3′; hence, we ensure a 1:1 donor-to-acceptor labeling stoichiometry. We prepared the corresponding donor-only reference sample by replacing the unmodified nucleotide Y(25) with G(25).

### Multiparameter Fluorescence Spectroscopy (MFS) for single-molecule FRET experiments (smFRET)

The home-built confocal system used for MFS-smFRET and calibration was described in detail elsewhere (30,38–40). To briefly summarize, we use Pulsed Interleaved Excitation (PIE) (41), or Alternating Laser EXcitation (ALEX) (42), with two diode lasers (Model LDH-D-C- 485 at 485 nm and laser LDH-D-C-640 at 640 nm; PicoQuant, Germany) operating at 40 MHz with 25 ns interleaved time. The power at the focal point of the 60X Olympus objective was set to 60 μW for the 485 nm laser line and 23 μW for the 640 nm excitation. The emitted photons were collected from the same objective and then spatially filtered by a 70 μm pinhole. The signal was separated into parallel and perpendicular polarizations with respect to the polarization of the excitation source, then split further into two different spectral windows defined as “green” and “red.” We used four photon-counting detectors, two for the green (PMA Hybrid model 40 PicoQuant, Germany) and two for the red (PMA Hybrid model 50, PicoQuant, Germany) channels. A time-correlated single-photon counting (TCSPC) module (HydraHarp 400, PicoQuant, Germany) in Time-Tagged Time-Resolved (TTTR) mode was used for data registration.

For smFRET experiments, samples were diluted to pM concentration in one of four buffers (*apo buffer:* 20 mM Tris and 200 mM NaCl at pH 7.0; *Mg^2+^ buffer:* 20 mM Tris, 200 mM NaCl, and 1 M MgCl_2_ at pH 7.0; *TPP buffer:* 20 mM Tris, 200 mM NaCl, and 2 mg/mL TPP at pH 7.0; *Mg^2+^ & TPP buffer:* 20 mM Tris, 200 mM NaCl, 1 M MgCl_2_, and 2 mg/mL TPP at pH 7.0). The buffers were charcoal-filtered to remove residual impurities. The concentrations of TPP and Mg^2+^ in these solutions are high to ensure saturating conditions. The particularly high concentration of Mg^2+^ is well beyond the physiological concentration of approximately 1-3 mM. This was used to demonstrate the extreme concentration necessary to stabilize a population of the aptamer domains in the closed configuration (see Results, Figure 3). At pM concentrations of the riboswitch, we observed ~2 molecules per second in the detection volume. NUNC chambers (Lab-Tek, Thermo Scientific, Germany) were pre-treated with a solution of 0.1% Tween 20 (Thermo Scientific) in water for 30 min and then rinsed with ddH_2_O to prevent adsorption artifacts. Data collection times varied from several minutes up to 10 hours. In long-time measurements, an oil immersion liquid with an index of refraction matching that of water (Carl Zeiss, Germany) was used to prevent drying out of the immersion water. Control experiments consisted of measuring distilled water to characterize the instrument response function (IRF), buffers for background subtraction, and nM concentration green and red standard dyes (Rhodamine 110, Rhodamine 101, and Alexa 647) in water solutions for calibration of green and red channels.

The data analysis used software developed in the Seidel group and is available at http://www.mpc.hhu.de/en/software (37,38,43–45). MFS accounts for many challenges associated with smFRET experiments, including photo-bleaching, cis-trans acceptor blinking, and dynamic and static quenching. We used FRET indicators such as the ratio of donor to acceptor fluorescence (F_D_/F_A_) and the average fluorescence lifetime of the donor (⟨τ_*D*(*A*)_⟩_*f*_), integrated over single-molecule events, or bursts (43,46,47). Details on the correction parameters obtained during calibration can be found in Supplemental Information Table S2. Bursts were selected for analysis based on the relative fraction of photons observed from the donor vs. the donor plus the acceptor fluorescence signals following their respective direct excitation pulses (*S_PIE_*) in order to ensure 1:1 active donor-to-acceptor stoichiometry (SI Figure S2, SI Methods).

### Time-Correlated Single Photon Counting (TCSPC) and filtered Fluorescence Correlation Spectroscopy (fFCS)

To identify stable populations that persist longer than the duration of the fluorescence lifetime of the FRET-labeled riboswitch (ns timescales), we compared the donor-only reference riboswitch with the FRET-labeled riboswitch in the same buffer conditions using TCSPC. Time-resolved fluorescence decay curves from photon arrival-time histograms (SI Figure S3, henceforth fluorescence decay histograms) ensure enough statistical rigor through large photon counts (on average >10^9^ photons for our FRET-labeled samples). Therefore, we determined the interdye distance distributions from fluorescence decay data by considering the donor-only and FRET-labeled riboswitch at saturated conditions of Mg^2+^, TPP, and Mg^2+^ and TPP, with the corresponding control in the absence of ligand (apo).

To arrive at the correct number of conformational ensembles, we fit the fluorescence decay histograms with an increasing number of fluorescence decay lifetimes corresponding to Gaussian-distributed FRET distances. Gaussian-distributed distances provide a physical representation of the mobility of the fluorescent labels and intrinsic dynamics of the chain. Although the shape of the distribution is less important, the mean and the variance are both relevant statistical parameters in single-molecule FRET experiments (48). Therefore, we fixed the variance according to benchmark studies (37–39,44) and optimized for the mean. Next, we evaluated the best results by visual inspection of the weighted residuals, monitoring the distribution of the autocorrelation of the residuals, and by a statistical improvement of the figure of merit, *χ*^2^ (31,49) (SI Equation 7). The model of the fluorescence decay includes a No-FRET population that accounts for inactive acceptor molecules (50) and for the extended conformations observed in the multidimensional histograms beyond what could be resolved by FRET. As expected, with the addition of distributions the figure of merit, *χ*^2^, and quality of the fit improves. We selected a model of two Gaussian distributions because the change of *χ*^2^ improved from an average of 8.52 over all conditions when using a single Gaussian distribution to 2.42 when using two Gaussian distributed FRET distances. Adding a third Gaussian distribution only marginally improved the average *χ*^2^ to 2.05 (SI Table S3).

To increase the photon counts and improve statistics for data analysis, TCSPC and *f*FCS measurements were performed using the MFS setup described above, but the power at objective was set for 120 μW for the 485 nm laser line and 39 μW for the 640 nm excitation while labeled samples were measured at nM concentration. The durations of these experiments were 10 hours each.

TCSPC and *f*FCS use the same dataset but consist of different analysis. TCSPC-generated fluorescence decays were jointly analyzed, along with a reference sample, to identify conformational ensembles that are stable in the nanosecond timescale with high precision. *f*FCS, a variation of standard FCS (51,52) that probes changes in the fluorescence lifetime (53), spectral information, and anisotropy (33), was used to probe transitions between selected conformers or pseudo-conformers characterized by specific fluorescence decay patterns. Thus, it was possible to identify changes in FRET due to conformational changes that occurred at timescales faster than the diffusion time. Details of the functional forms used to process the data, the statistical tests, and error analysis are found in the Supplemental Information.

For fFCS, we systematically tested models of increasing complexity through the addition of relaxation times (t_R_) and finally chose a model by iteratively varying the number of terms in each factor in the *f*FCS function to find the number of terms necessary to adequately fit the data, as determined by visual inspection of the residuals, and to provide a significant relative improvement in the *χ*^2^ compared to the use of one fewer term. The global fit model we used here has four distinct relaxation terms describing conformational transitions and two exponential terms to account for dark state kinetics (in the case of acceptor, this can be due to cis-trans isomerization, and in the case of the donor, this represents a transition to triplet state) and photobleaching (33).

### Discrete Molecular Dynamics (DMD) simulations

DMD is an event-driven approach to molecular dynamics (MD) simulations that utilizes discontinuous step-functions instead of continuous functions to model the interaction potentials between atoms (54). Compared to traditional MD simulations that calculate the forces and accelerations of all atoms, DMD computes the ballistic equation of motion, considering conservation laws of energy, momentum, and angular momentum, for those atoms involved in a collision event, defined as two interacting atoms within the distances associated with potential steps, allowing longer length and time scales in simulating complex biomolecular systems (35,55).

The coarse-grained RNA model uses three beads corresponding to the phosphate, sugar, and base groups of each nucleotide (36). The interaction potentials include base paring, stacking, and the effective loop energy, which were tabulated from experimental measurements (56). DMD simulations with the coarse-grained RNA model can successfully predict RNA 3D structures and folding kinetics (57,58).

The atomistic RNA model adopts the united-atom approach to model nucleotides, which includes all heavy atoms and polar hydrogen atoms. Similar to all-atom DMD simulations of proteins (35), bonded interactions, including covalent bonds, bond angles, and dihedrals, were assigned according to a statistical analysis of high-resolution RNA structures in the protein databank (PDB). We used the Medusa force field (57) to describe the non-bonded interactions, including van der Waals (VDW), hydrogen bonding, solvation, and electrostatics. The VDW parameters were taken from the CHARMM19 united-atom force field, and the distance and angular dependent hydrogen bonds were explicitly modeled by a reaction-like algorithm (59). The screened electrostatic interactions between charged atoms were computed using the Debye-Hückel approximation, while the solvation term for implicit solvent simulations was modeled by the Lazaridis-Karplus EEF1 (Effective Energy Function 1) model (60).

The initial structure of TPP-riboswitch was obtained from the protein data bank (PDBID: 2CKY (17)). Periodic boundary conditions were imposed in all simulations, and the riboswitch was initially positioned in a cubic box with an edge length of 100 Å. Counter ions were added to maintain the overall charge neutrality of the system. Table S1 of Supplemental Information lists the number of Mg^2+^ and Na^+^ ions in the simulated systems. Production runs followed 2000 timestep energy minimization runs, where a DMD timestep corresponded to ~50fs. To efficiently sample the conformational space of the free energy landscape as well as the dynamics of riboswitch closing pathways, we employed replica-exchange DMD (REXDMD) simulations in our study. In the replica-exchange scheme, 16 replicas in consecutive temperature ranges within the total range 260-335 K were simulated via the exchange of temperature between replicas at periodic time intervals, thus overcoming any local energy barriers during the conformational sampling of the energy landscape. The simulation temperature was maintained using the Andersen thermostat (61). Each replica lasted 4 × 10^6^ time steps, or approximately 200 ns of simulation time totaling ~3.2 μs of simulation time.

Since the smFRET experiments focused on the open and close conformational dynamics of two stems in the TPP riboswitch and the RNA was expected to maintain the native-like base pairing at the observed fast time-scales, we imposed base-pair constraints in our all-atom REXDMD simulations to decouple the three-dimensional conformational dynamics of tertiary structures from the folding and unfolding dynamics of helices that occur on longer timescales (62). Similarly, we also assumed that TPP stayed in its native binding pocket by imposing distance constraints between TPP and two stacking nucleotides. For interactions of Mg^2+^ with RNA, we only considered VDW, solvation, and electrostatics without including the effect of metal coordination.

### Weighted histogram analysis method (WHAM)

The thermodynamics of TPP riboswitch conformational dynamics were computed with WHAM analysis (63,64). The density of states, *g*(*E*), was calculated self-consistently by combining the energy histograms from all replicas, using the last 100 ns of replica-exchange trajectories. The potential of mean force (PMF), i.e., the effective free-energy landscape, was computed as a function of a given physical parameter, *x*:

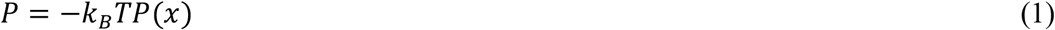

where *k*_B_ is the Boltzmann constant, *T* denotes the temperature of interest, and *P*(*x*) corresponds to the probability of finding the riboswitch with specific values of *x*. *P*(*x*) was computed according to:

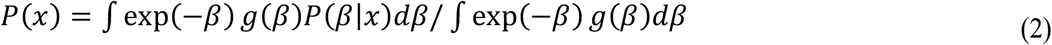

where *β*=1/*K_B_T* and *P(E*|*x)* is the conditional probability of finding conformations with parameter value *x* while the energy is in the interval (*E, E+dE*). The *x* parameter can be comprised of multiple variables. For example, we evaluated the conformational free energy landscape of TPP riboswitch with respect to both the inter-arm distance, *D*_inter-arm_, as measured between the 2′-hydroxyls of labeled G25 and U56, and with respect to the P1/P2 co-stacking distance, *D*_*co*-stack_, as measured between U39 in P2 and G73 in P1.

### Clustering analysis

The hierarchical clustering program (www.compbio.dundee.ac.uk/downloads/oc) was used to group similar conformations of the riboswitch. Depending on an input pair-wise root-mean-square distance (RMSD) matrix, the clustering algorithm iteratively combined the two closest clusters into one. The “cluster distance” was calculated based on all pairwise distances between elements of two corresponding clusters. We used the mean of all values to compute the distance between two clusters, and the centroid structure of each cluster was chosen such that it had the smallest average distance to other elements in the cluster.

### RNA preparation for optical tweezer experiments

Optical tweezer experiments consisted of the TPP riboswitch aptamer domain flanked at both ends by single-stranded DNA extensions (sequence in the Supplemental Information) annealed to 2 kbp DNA handles, each with a single-strand overhang. The DNA handles were amplified from an arbitrarily selected stretch of the pMAL-c5X vector (N8108, New England BioLabs) with primer sequences detailed in the Supplemental Information. The 5’ end of the forward primer of Handle 1 was modified with an amine group (NH_2_) for binding to the N-hydroxylsuccinimide (NHS) group of N-hydroxylsuccinimide functionalized polyethylene glycol (PEG-NHS) on the coverslip. To ensure a single strand overhang, we used an autosticky reverse primer (65) with the tetrahydrofuran abasic site inserted 35 bases from its 5′ end, blocking the synthesis of the complementary strand for a 35-nucleotide 5′ overhang that anneals to the 5′ DNA extension of the RNA construct. The 5′ end of the forward primer for Handle 2 was modified with biotin for linking to streptavidin-coated beads. Three phosphorothioate bonds were inserted 37 nucleotide bases from the 5’ end of the forward primer, inhibiting nuclease digestion (66). Handle 2 was then digested with Lambda Exonuclease (M0262, New England BioLabs), which removes nucleotides from linearized double-stranded DNA in the 5′ to 3′ direction, leaving a 3’ overhang on the complementary strand that anneals to the 3’ DNA extension of the RNA construct (67).

The efficiency of RNA construct to DNA handle annealing was determined empirically by observing which dilutions of the constructs gave enough tethers to beads during tweezers experiments (68). To minimize the likelihood of finding RNA constructs without both handles, we annealed the handles with the RNA construct in 1:0.25:1 (H1:RNA:H2) molar ratio in PBS (pH 8.3) buffer at room temperature for 2 hours.

Flow cells (~10 μL) were made by cutting a channel (2-3 mm wide) into parafilm sandwiched between a glass slide and 22 × 22 mm cover glass, pre-cleaned with piranha solution (sulfuric acid and hydrogen peroxide in the ratio of 3:1) followed by series of ethanol and distilled deionized water washing steps (69). The piranha solution, being a strong oxidizing agent, cleaned off most of the organic residues and hydroxylates from the surface to facilitate the subsequent covalent bonding with silanes.

We conjugated the RNA construct to coverslip surface by adapting a standard protocol (70). Briefly, a 2% (w/v) silane-PEG-NHS (Nanocs, NY) solution prepared in pure, dry DMSO was flowed into the flow cell and incubated at room temperature for 2 hours. The annealed handle-RNA construct with -NH_2_ and biotin labels at either end was then flowed into the flow cell and incubated in a humidity chamber for 1.5 hours to allow covalent bonding between the NHS and the amine modification on DNA handle. 1.1% (w/v) 1.05 μm diameter streptavidin-coated polystyrene beads (Bangs Lab) were washed twice in one of the buffers detailed above (i.e., apo buffer, TPP buffer, Mg^2+^ buffer, or Mg^2+^ & TPP buffer) plus 0.1% Tween 20. The bead solution was then diluted 20 times and flowed into the flow cell. The flow cell was then rinsed with 1% Pluronic F-127, and the ends were sealed with silicone grease. The flow channel was left to sit at room temperature for 20-24 hours.

### Optical tweezers and data acquisition

A custom-built, single-beam optical tweezer with a 1064 nm, ytterbium fiber laser (YLR-10-1064-LP, IPG Photonics) focused into the sample plane by a CFI60 Plan Apochromat Lambda 60x N.A. 1.4 Oil Immersion Objective Lens (Nikon) was used to trap the beads and pull on the handles of the riboswitch. Measurements of the change in the construct length resulting from the pulling were made by determining the displacement of the trapped bead from the optical trap center using quadrant photodiode (QPD) back focal plane detection (71) as well as the translation of the nanostage (Nano-LP100, Mad City Labs Inc.) holding the cover glass. Force measurements were made by obtaining calibration parameters (72), detection sensitivity (β in V/μm) and stiffness (κ in pN/nm), from a non-linear Lorentzian fit of the power spectrum density of the QPD response to a trapped bead subject to thermal fluctuations. The parameters were used to convert the change in QPD voltage signal due to displacement of trapped bead into a corresponding distance (nm) and then that displacement to a force (pN). A trap stiffness of 0.2-0.3 pN/nm was used for all measurements. The data were acquired using an FPGA (PXI-7854R, National Instruments) and custom-written LabView VIs (National Instruments).

The passive force time-series data were recorded at 1000 Hz for 100 s at each force and filtered offline using a 25 ms moving average filter (MATLAB, MathWorks). Under the passive mode force setting, the optical tweezer was used with a fixed trap where both the force and extension vary as the molecule undergoes structural transitions.

The MATLAB function findchangepts was used to partition each of the 100 s time series data into various force levels. The findchangepts function minimizes the sum of the residual error of a particular region in the dataset from its local mean and returns a step at which the mean changes the most. Depending upon the position of the stage, and hence the average force-extension of the RNA construct, the obtained mean forces were fitted to 2 or 3 Gaussian functions (using the MATLAB Gaussian mixture model). The transition forces determined from the multiple-Gaussian fit were used to define the folded (denoted by higher force or shorter extension) and unfolded (denoted by lower force or longer extensions) states. The mean dwell time of a folded state was obtained by fitting the dwell times distribution of the state to a single decay exponential function and calculating the characteristic dwell time constant from the fit. The corresponding rate coefficient (unfolding rate, *k*_on_) was calculated as the inverse of the mean dwell time. To first order approximations, this rate constant depends exponentially upon the applied force as given by two parameters Bell’s equation, (73)

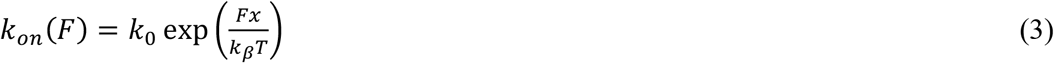

where *F*is the applied force, *k_0_* is the rate at zero force, and *k_β_* is Boltzmann’s constant, *T* is the absolute temperature, and *x* is the characteristic distance the stretch. The logarithm of *k*_on_ plotted as a function of force was fitted with a linear equation to determine the *k*_on_ at zero force from the y- intercept.

For force extension experiments, the stage was repeatedly translated along a horizontal direction at a step size of 5 nm. At each position, after a wait time of 15 ms to a steady-state condition, data were collected for 2 ms at a 500 kHz rate and averaged before moving to the next position. The unfolding rips were identified by sudden drops in force, which correspond to local maxima in the force extension curve (FEC).

## RESULTS

### Sub-millisecond dynamics of Arabidopsis thaliana TPP riboswitch’s aptamer domain

We placed the FRET-labeled aptamer domain of the TPP riboswitch (referred to as the TPP riboswitch) in various buffer conditions, and we observed changes in the shape and position of the two-dimensional MFS histograms (Figure 2). In the apo buffer (absence of Mg^2+^ and TPP), we observed a narrow, unimodal distribution centered in the low-FRET regime (Figure 2A). Even at extreme saturated Mg^2+^ concentrations (>100 mM, Figure 2B), the population was unimodal and centered in the low-FRET regime, with a slight decrease in the mean and increase in the skew of the fluorescence-weighted lifetime population as compared to the apo ^buffer results^ according to MFS alone. The population still lies along the static FRET line.

**Figure 2.**
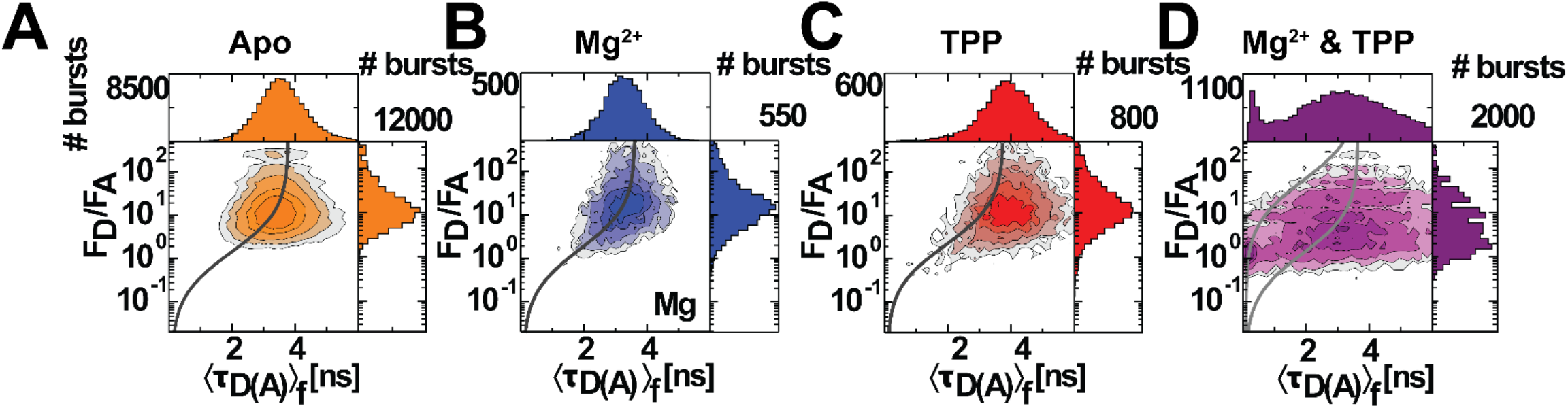
The dynamical nature of the TPP riboswitch aptamer domain depends on the presence of TPP and Mg^2+^. Multidimensional smFRET populations for the TPP riboswitch in A) Apo, B) Mg^2+^, C) TPP, and D) Mg^2+^&TPP buffers (see Methods for buffer compositions; total number of single-molecule events, bursts, are N=94,999, N=5,198, N=8418, and N=21,836, respectively). The FRET indicators shown are the ratio of donor to acceptor fluorescence (F_D_/F_A_) and the average donor fluorescence lifetime (⟨τ_*D*(*A*)_⟩_*f*_) for each PIE-selected 1:1 donor-to-acceptor stoichiometry burst (see Methods). Darker contours correspond to a higher density of bursts, and static FRET lines (black line, SI Table 2) describe the expected relationship between the FRET indicators for non-exchanging, static FRET populations according to Förster theory. Large F_D_/F_A_ and large ⟨τ_*D*(*A*)_⟩_*f*_ correspond to the low FRET regime (long interdye distance), and small F_D_/F_A_ and ⟨τ_*D*(*A*)_⟩_*f*_ correspond to the high FRET regime (short interdye distance). Note that two static FRET lines are needed in Mg^2+^&TPP conditions (panel D) due to acceptor quenching (see Supplemental Information). Further, it is qualitatively clear that the addition of TPP or Mg^2+^ & TPP shifted the observed populations toward high FRET and skewed to the right of the static FRET line, as is expected for dynamically exchanging FRET populations.

**Figure 3.**
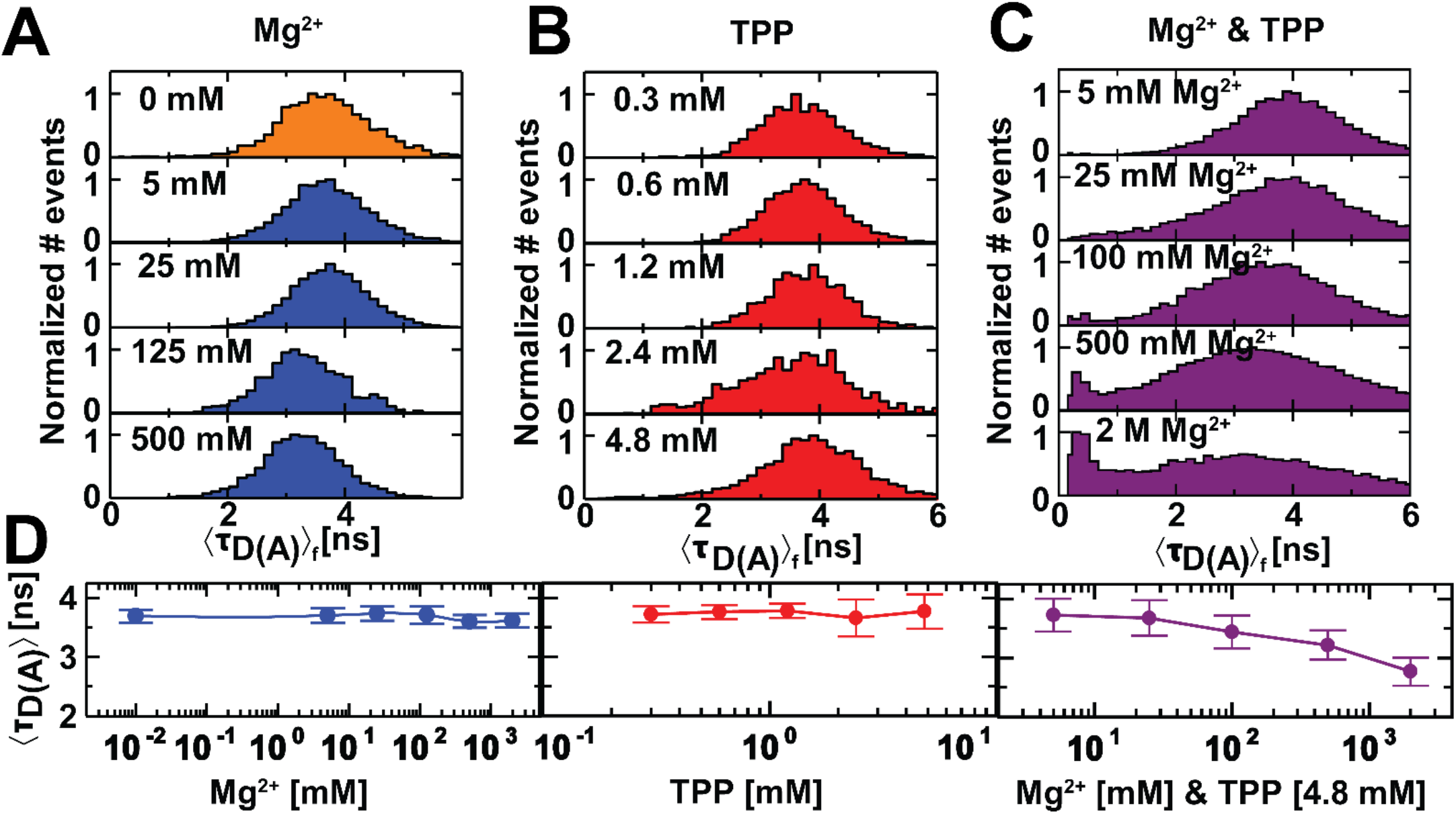
Increasing concentrations of Mg^2+^ in presence of TPP induce a population of closed states in the aptamer domain. Mean donor fluorescence lifetime distributions for FRET-labeled TPP riboswitch (A) with increasing concentrations of MgCl_2_ starting from 0 mM to 0.5 M, (B) with increasing concentrations of TPP starting from 0.3 mM to 4.8 mM, (C) with increasing concentrations of MgCl_2_ starting from 5 mM to 2 M, all at a fixed TPP concentration of 4.8 mM. (D) Mean donor fluorescence lifetimes calculated from the data in (A), (B), and (C). The standard deviations of the distributions are shown as error bars.

In the TPP buffer, we found a broader distribution of the FRET measurement population that tailed toward lower F_D_/F_A_ values (Figure 2C), which indicates the presence of a more closed, higher-FRET states that we did not observe in either the apo or even at extreme Mg^2+^ concentrations (Figure 2A-B). Additionally, in the TPP buffer, we found that the highest density of FRET measurements was shifted slightly to the right of the static FRET line (Figure 2C), which indicates dynamic averaging of FRET states within the ms measurement timescale (30,43,74) that was not observed in the apo or Mg^2+^ buffers (Figure 2A-B). Together, these results suggest that, in the presence of TPP alone, the riboswitch’s aptamer domain exhibits conformational dynamics between open, low-FRET states and a more closed, higher-FRET states at timescales faster than the millisecond observation time.

In the Mg^2+^ & TPP buffer, we found two distinct FRET populations, which was particularly evident in the one-dimensional projection of the ⟨τ_*D*(*A*)_⟩_*f*_ distribution (Figure 2D). One of these populations was broad (~3.5 ns FWHM) and centered around ⟨τ_*D*(*A*)_⟩_*f*_ = 3.1 ns (medium-FRET), and the other was narrow (~0.3 ns FWHM) and centered below ⟨τ_*D*(*A*)_⟩_*f*_ < 0.5 ns (high-FRET) (Figure 2D). This splitting of the FRET populations (Figure 2D), as compared to the results in other buffers (Figure 2A-C), suggests that the exchange processes between open and closed configurations occur at timescales similar to or longer than ms timescale (reported for each sample, Table S4E) (30,74) in either saturating Mg^2+^ or TPP buffer conditions. This result suggests only the presence of both saturating TPP and extreme Mg^2+^ concentrations slows transitions away from the high-FRET closed state of the aptamer domain to a timescale greater than the ms. In any other condition (Figure 2A-C), there is not a measurable long-lived (>ms) population of closed aptamer domains. Additionally, the broadening of the lower-FRET population in TPP and shift of its peak to medium-FRET upon the addition of Mg^2+^ suggests that Mg^2+^ may enable the aptamer domain to sample other, intermediate states along the closing pathway.

Together, these results suggest that the *Arabidopsis thaliana* TPP riboswitch’s aptamer domain is in an open state (with L3 and L5 loops far apart, Figure 2A-B) in the absence of TPP, can transiently be found in a more closed state with TPP alone (with L3 and L5 loops closer together, Figure 2C), and populates a long-lived (>ms timescale) closed state (predicted to be the L3 and L5 loops close together like the Mg^2+^ & TPP bound structural model) in only the presence of saturating TPP and Mg^2+^ concentrations (Figure 2D) (17). Our results also indicate that the aptamer domain responds to its two ligands, Mg^2+^ and TPP, differently (Figure 2B and C), which suggests two independent closing pathways. These results do not conclusively determine whether the aptamer domain is dynamic in the apo or Mg^2+^ alone conditions (Figure 2A and B), as opposed to the TPP alone condition, in which we observed a shift away from the static FRET line (Figure 2C). Nonetheless, the observed differences in the shape of the FRET populations (Figure 2) suggest a dynamic state of the aptamer domain in all measured conditions (30,43). To probe the dynamics of the TPP riboswitch’s aptamer domain further we use TCSPC and *f*FCS, detailed in the following sections.

### The stabilizing effect of extreme Mg^2+^ concentrations on the aptamer domain

To further address the role of Mg^2+^ ions ability to coordinate conformational changes in the aptamer domain, we measured the donor fluorescence lifetime at increasing concentrations of Mg^2+^ (0 - 0.5 M, Figure 3A) in the absence of TPP. We found that the mean donor fluorescence lifetime (⟨τ_*D*(*A*)_⟩_*f*_) population distributions were nearly invariant over this broad range of Mg^2+^ concentrations (Figure 3A). In particular, we found that the expected values and the standard deviations of the mean donor fluorescence lifetime population distributions were not significantly different over this range of Mg^2+^ concentrations (Figure 3D, left). This result suggests that the TPP riboswitch’s aptamer domain stays open in the absence of TPP, even for increasing Mg^2+^ over a large, non-physiological range of concentrations.

We measured the mean donor fluorescence lifetime of the FRET-labeled TPP riboswitch at increasing concentrations of TPP (0 - 4.8 mM, Figure 3B), but in the absence of Mg^2+^, and found that the donor fluorescence lifetime population broadened with increasing TPP concentration. In particular, we found that the expected values of the mean fluorescence lifetimes were relatively constant while the standard deviations of the mean donor fluorescence lifetime population distributions increased concomitantly with TPP concentration (Figure 3D, middle). This broadening was consistent with our previously noted observation of a broader mean donor fluorescence lifetime in the TPP buffer as compared to the apo buffer (Figure 2C and A, respectively).

Because we had found that the FRET population deviated from the static FRET line in the TPP buffer (Figure 2C), which is consistent with dynamic averaging effects and indicates that the riboswitch’s aptamer domain is unable to be stabilized in the closed state in the absence of Mg^2+^, we investigated the effect of Mg^2+^ concentration on the ability of the aptamer domain to be in the high-FRET closed state by measuring the donor fluorescence lifetime of the FRET-labeled riboswitch at increasing concentrations of Mg^2+^ in TPP-saturating (4.8 mM) conditions. We found that at a 5 mM concentration of Mg^2+^, which already is notably high compared to the expected physiological Mg^2+^ concentration of about 1 mM in *Arabidopsis thaliana* (75), the mean donor fluorescence lifetime distribution of the TPP riboswitch’s aptamer domain (Figure 3C, top) is not significantly different from the case of saturating TPP and no magnesium (Figure 3B, bottom). When the concentration of Mg^2+^ is increased to 25 mM, we found a broadening of the mean donor fluorescence lifetime distribution due to an increase in the frequency of observations at the higher-FRET tail (Figure 3C). It is only as the Mg^2+^ concentration is further increased to extreme limits (to 100, 500, and 2000 mM, Figure 3C), that we find a significant high-FRET population, indicated by the population peak at lower ⟨τ_*D*(*A*)_⟩_*f*_ values (Figure 3C), a corresponding lower expected values (Figure 3D, right), and an increase in the standard deviation of the total population (Figure 3D, right). Together, these results suggest that Mg^2+^ stabilizes a closed state of the aptamer domain only at non-physiological concentrations of Mg^2+^, which we expect is most likely due to the screening effect of Mg^2+^ against the repulsive negative charges on the RNA backbone and the electronegative polarity of TPP. In total, these results strongly suggest that the TPP riboswitch aptamer domain exists in a highly dynamic state that transiently samples closed-like configurations at physiological concentrations of magnesium.

### There are at least five distinct ensembles of the aptamer domain

To identify conformational ensembles of the aptamer domain that persist longer than the duration of the fluorescence lifetime of the FRET-labeled riboswitch (ns timescales), we compared the fluorescence decay histograms of donor-only reference riboswitches, which we generated using time-correlated single-photon counting (TCSPC, Methods), with the decay histogram of the FRET-labeled riboswitch in the same buffer conditions. We found that there was a significant population of conformational ensembles with no FRET (gray bars) compared to the corresponding fraction with donor fluorescence quenched through FRET (colored bars) in each buffer condition (Figure 4A). The fraction of molecules exhibiting FRET increased from 27% in the apo buffer to 41% in saturating Mg^2+^ & TPP conditions (Figure 4A). This result is consistent with the multidimensional smFRET data, in which we observed a population shift toward high-FRET (Figure 2) and a broadening of the ⟨τ_*D*(*A*)_⟩_*f*_ distribution (Figure 3) in Mg^2+^ & TPP buffer as compared to apo buffer.

**Figure 4.**
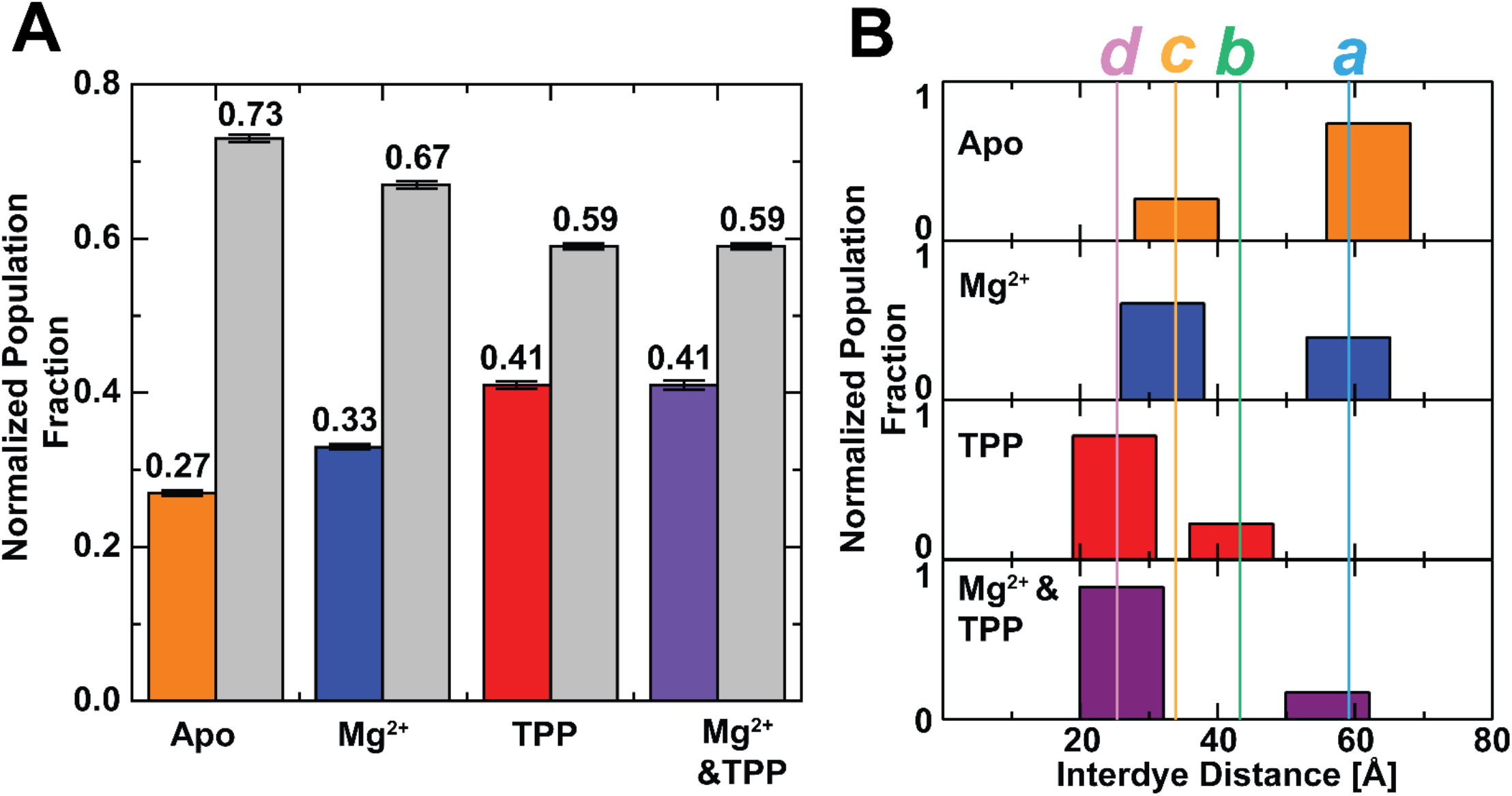
Identification of at least five aptamer domain conformational states as a function of ligand concentration. (A) Bar plot represents the fraction of the population of molecules exhibiting FRET (color-filled bars) and no-FRET (gray bars) as determined by fluorescence decay from TCSPC FRET experiments. B) Interdye distance distribution with a two-Gaussian distribution fitting model. State *a* is semi-open state, state *b* is the intermediate state I, state *c* is the intermediate state II, and state *d* is the closed state of the aptamer domain.

We calculated the interdye distances of the FRET-exhibiting population of TPP riboswitches in each buffer using the functional model (SI Equation 4). We identified a total of four distinct TPP riboswitch aptamer domain ensembles (*a, b, c*, and *d*, Figure 4B), with at least two at each buffer condition undergoing dynamic interconversion between those ensembles. In each of the buffers that contain ligand (TPP, Mg^2+^, or both) the more populated FRET-exhibiting state was the more closed state (smaller mean interdye distance, ⟨*R*^(1)^⟩, Figure 4B), while in the apo buffer the more open state (larger mean interdye distance, ⟨*R*^(2)^⟩, Figure 4B) was more populated. Further, in the ligand-containing buffers, the no-FRET population was smaller compared to the apo buffer. In the apo and Mg^2+^ buffers, we found that the FRET-exhibiting states ⟨*R*_DA_⟩s were similar to one another (states *a* and *c*, Figure 4B and Table 1), but the relative population fractions switched, with the more closed state *c* being more populated in Mg^2+^ and the more open state *a* being more populated in apo buffer (Figure 4B). The change in relative state populations suggests a change in the exchange dynamics between these ensembles but does not reveal whether this is due to faster transitions from *a* to *c* or slower transitions from *c* to *a*, as the MFS histograms appear unimodal for both conditions. Thus, the difference is likely due to subtle changes in the sub-ms regime. In both buffers containing saturating concentrations of TPP (TPP and Mg^2+^ & TPP buffers), where the fraction of the population exhibiting FRET was 41% (Figure 4A), we found that 78% of the FRET-exhibiting ensemble had a mean interdye distance of 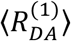 = 25.4±1.0 Å (state *d*, Figure 4B). The other 22% of the FRET-exhibiting ensemble had 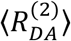 =43.4±1.2 Å in TPP alone (state *b*, Figure 4B) and 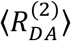 = 62.1±4.5 Å (state *a*, Figure 4B) in the Mg^2+^ & TPP buffer. We also found that the most-populated FRET-exhibiting ensemble observed in both TPP-containing buffers 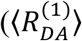 = 25.4±1.0 Å, state *d*, Figure 4B) is more closely related to the most-populated FRET-exhibiting ensemble observed in the Mg^2+^ and apo buffers (interdye distance of 31.7±1.9 Å, state *c*, Figure 4B) than the less-populated ensembles in TPP-containing buffers (states *a* and *b*). Finally, we found that in the Mg^2+^ & TPP buffer the more open state was like state *a* found in the apo and Mg^2+^ buffers rather than state *b* found in the TPP buffer (Figure 4B). A summary of the interdye distances and their statistical uncertainties are shown in Table 1. We found that ensembles *d* and *c* (Figure 4B) agreed within 5 Å with Accessible Volume (AV) modeling of the dyes at the corresponding locations in a structural model thought to be associated with the TPP riboswitch OFF state (26.9 Å, SI Figure S4) (37).

**Table 1.** Interdye distances associated with the conformational states of the aptamer domain identified in the population of FRET-exhibiting TPP-riboswitches. Population fractions of the two Gaussian distributed states are normalized, such that *x*(1) + *x*(2) = 1. The statistical uncertainties, ε, for the mean interdye distances are estimated using both the widths of the distributions and uncertainty due to dye reorientation (described in SI). The upper index in parenthesis is the corresponding numbered state. Distances are color-coded according to the state assignments in Figure 4B, state *a* is blue, state *b* is green, state *c* is orange, and state *d* is red.

| Sample | $\langle R_{DA}^{(1)} \rangle \pm \varepsilon$ [Å] | $x^{(1)}$ | $\langle R_{DA}^{(2)} \rangle \pm \varepsilon$ [Å] | $x^{(2)}$ |
| --- | --- | --- | --- | --- |
| Apo | $34.0 \pm 2.3$ | 0.26 | $62.1 \pm 4.5$ | 0.74 |
| Mg<sup>2+</sup> | $31.7 \pm 1.9$ | 0.61 | $58.7 \pm 3.3$ | 0.39 |
| TPP | $25.4 \pm 1.0$ | 0.78 | $43.4 \pm 1.2$ | 0.22 |
| Mg<sup>2+</sup> & TPP | $26.1 \pm 1.2$ | 0.83 | $55.6 \pm 2.5$ | 0.17 |

Our data suggest the existence of aptamer domain ensembles in which the fluorescent labels are too far apart distance to be quantified by FRET due to low acceptor sensitization. Therefore, TPP riboswitch aptamer domains in this state contributed to the fraction of the population in the no-FRET, open, state (interdye distance >80 Å, Figure 4A, grey bars). We have confidence in the presence of these long interdye distance populations because the synthesis of the FRET-labeled riboswitch ensures 1:1 donor-to-acceptor stoichiometry (50), with analysis limited to bursts with *S_PIE_* values near 0.5 (SI Figure S2). However, this fraction may also include riboswitches that are not entirely folded as well as a small number that might contain acceptor fluorophores incapable of FRET.

In total, our results strongly suggest that there are at least five distinct ensembles, but only three limiting are resolved in each buffer condition, with one state for which the interdye distance is too long to be quantified with the chosen FRET pair and assigned labeling locations. Sampling of multiple aptamer domain ensembles in each buffer suggests both a dynamic aptamer domain closing pathway and a rheostat-like tuning of the aptamer domain configuration states that might control switching in the switching sequence between ON-like and OFF-like structural ensembles.

### The aptamer domain ensemble switching rates span four orders of magnitude

We identified at least five distinct aptamer domain ensembles (Figure 4 and Table 1), but the multidimensional smFRET histograms displayed mostly unimodal distributions with varying widths (Figures 2 and 3). To obtain a holistic solution, we postulate that the unimodal distribution in Figures 2 and 3, must reflect dynamic averaging occurring between 25 ns (TCSPC observation window) and the ms-timescale (the diffusion time of the TPP-riboswitch). To test this hypothesis, we used filtered fluorescence correlation spectroscopy *f*FCS), which, as opposed to traditional fluorescence spectroscopy methods, uses characteristic fluorescence decays and time-resolved anisotropy decays to probe state-specific exchange processes (33,34). When only two states undergo kinetic exchange, the cross-correlation function between species corresponding to those states shows a decrease in the amplitude of the correlation at the characteristic timescale of exchange between those states. Additional decay terms appear for additional transitions in more complex kinetic schemes. Neither species-specific auto-correlation function exhibits correlation amplitude decreases at these characteristic anticorrelation times (33,34). Thus, we use *f*FCS cross-correlation curves to probe the conformational transitions in the aptamer domain that are only reported by FRET changes.

We performed a global fit of the species-specific auto- and cross-correlation data (34) that simultaneously satisfied all the relaxation terms, all terms corresponding to photophysical effects, and all diffusion terms in the correlation functions (Methods, Figure 5) using the same photon-stream data as in the TCSPC experiments (Figure 4) for each buffer condition (SI Figure S4). We found relaxation times ranging from 100 ns to ms, with one observed at every decade in time (e.g., apo buffer, Figure 5A). We calculated the effective transition rate constants associated with each buffer condition by taking the inverse of the relaxation times (Figure 5A and 5B for apo buffer, SI Figure S5 for the other conditions) and plotting them for each measured condition (Figure 5C). These transition rates indicate switching between at least five (number of relaxation times plus 1) different configurations at timescales ranging from μs to ms, consistent with the assessment of number of ensembles derived by time-resolved analysis, even in apo conditions. Moreover, we found slightly slower effective rates at higher Mg^2+^ conditions compared to apo buffer conditions suggesting that the relative population changes observed in the donor fluorescence decays in these two conditions requires Mg^2+^. This mechanism would slow transitions from state *c* to state *a* at non physiological conditions. Further, increases in the effective transition rates in TPP and Mg^2+^ & TPP buffers indicate that a saturating concentration of TPP significantly lowers the energy barriers between the sampled ensembles and that more transition pathways are available between configurations.

**Figure 5.**
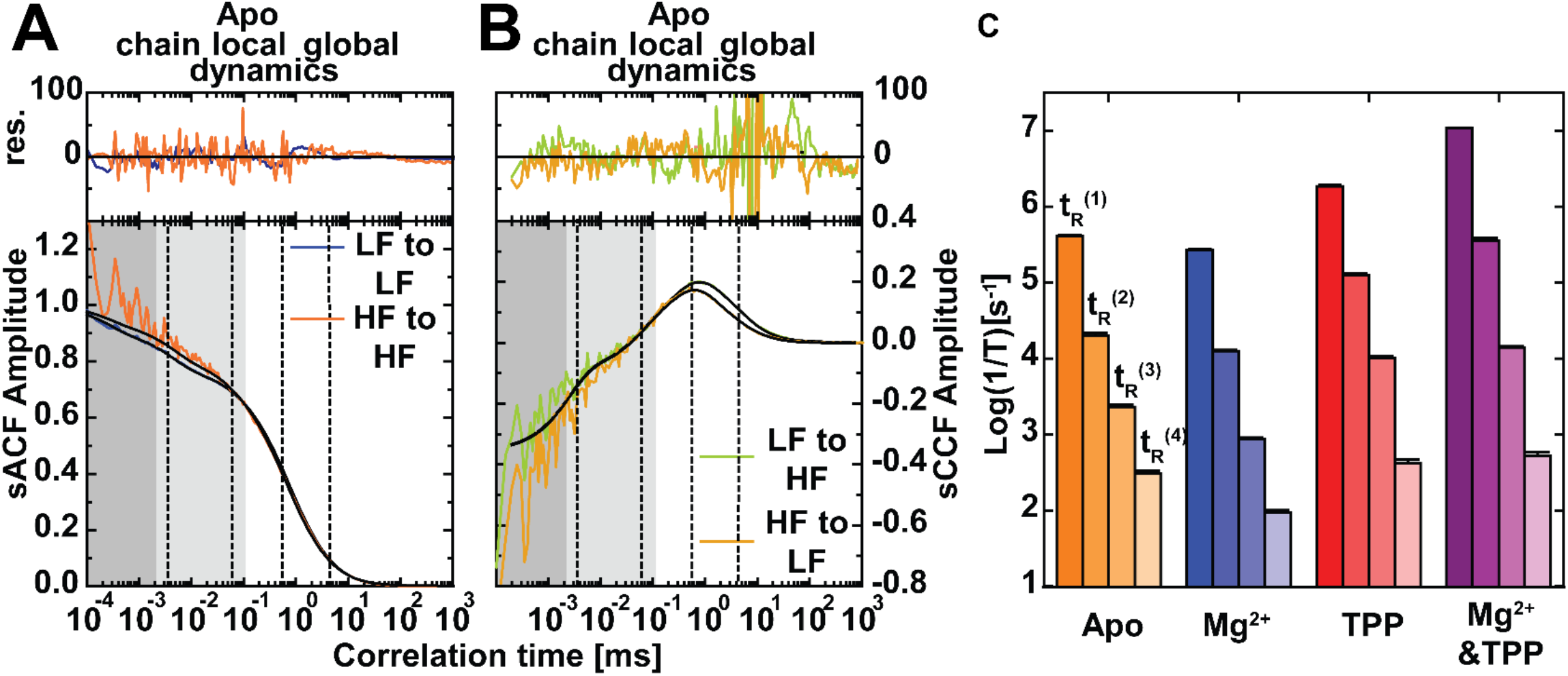
The aptamer domain exhibits four dynamic timescales. Example filtered FCS species auto (A, sACF) and cross-correlation (B, sCCF) functions from FRET experiments for the TPP riboswitch in the apo buffer. Four distinct state transitions rates (vertical dashed lines), spanning different decades in time were identified in all conditions, as exemplified here by data obtained in the apo buffer. Shaded regions correspond to timescales typical of chain dynamics, local conformational changes, and global dynamics (76,77), from darker to lighter, respectively. Functional fits are shown as solid black curves, while colored lines represent the raw correlation data. C) Four relaxation time populations were observed in each set of conditions. Bar height represents the log of the inverse of each correlation time. fFCS fit parameters are listed in Supplemental Information Table S4G.

### Ensemble switching observed with FRET is predicted by Replica Exchange DMD Simulations

To provide structural insights into the experimentally observed conformational dynamics, we employed replica-exchange discrete molecular dynamics (REXDMD) simulations of the riboswitch under different conditions, including in the absence and presence of TPP and/or Mg^2+^ ions (see Methods). We computed the probability distribution function (PDF) of the inter-arm distance between dye-conjugated bases G25 and U56 at room temperature (Figure 6A). In the absence of the TPP and Mg^2+^ co-ligands, we found that the TPP riboswitch’s aptamer domain stayed in open-like state conformations with the highest probability centered at an inter-arm distance of ~93.7 Å and a broad distribution tailing toward shorter inter-arm distances (Figure 6A). In the presence of Mg^2+^ at a 38:1 molecular ratio to RNA (~63.1 mM simulation concentration), the riboswitch exhibited a similar trend as in the apo form, and the PDF displayed a shoulder at the inter-arm distance of ~85.3 Å in addition to the peak centered at ~92.8 Å (Figure 6A). In the presence of TPP only (1:1, or ~1.6 mM), the riboswitch featured a PDF with a broad unimodal distribution centered at an inter-arm distance of 73.1 Å (Figure 6A). In the presence of both Mg^2+^ & TPP, the PDF had two distinct peaks, with the inter-arm distance centered at 24.8 Å representing a closed state ensemble and the population with peaks at 85.2 and 95.1 Å resembling the open state ensemble, similar to the apo and Mg^2+^ conditions (Figure 6A). These simulation results agreed well with the FRET measurements (Figure 4), which showed the appearance of low-populated closed-like ensemble (state *c*) in the Mg^2+^ buffer, a tailing toward a closed-like ensemble (state *d*) and the appearance of intermediate configurations (state *b*) in the TPP buffer, and a prominent population peak for a closed state (state *d*) and simultaneous reduction in the open-like ensemble (state *a*) in the Mg^2+^ & TPP buffer.

**Figure 6.**
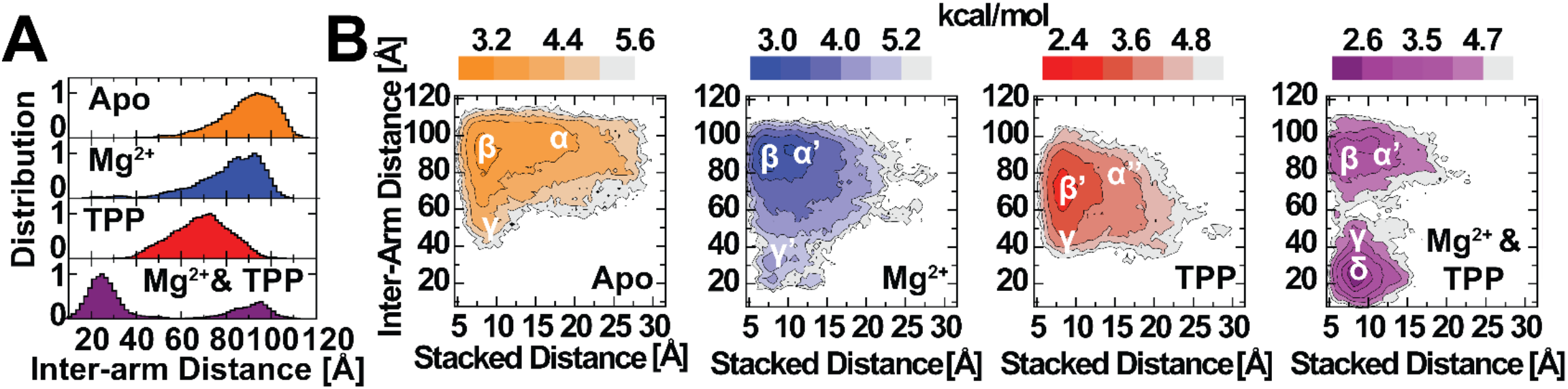
Atomistic DMD simulations of the riboswitch find aptamer domain conformational states that correlate to the FRET data. (A) The probability distribution function of the inter-arm distance between G25 and U56. (B) The two-dimensional PMF as a function of inter-arm (G25 and U56) and co-stacked distances (between U39 and G73). The basins correspond to open state with no co-stacking (α, α′, α″), open state with co-stacking (β, β’), partially closed state with co-stacking (γ, γ’), and closed state with co-stacking (δ).

The simulation trajectories and RNA conformations suggested that the co-stacking between stems P1 and P2 was coupled to the opening and closing dynamics of the arms. Hence, we computed the PMF (see Methods) as a function of both the inter-arm distance, *D_inter-arm_* (measured between the oxygen atoms of 2′-hydrogens of the labeled G25 and U56 and representing aptamer domain configuration), and the co-stacking distance, *D_co-stack_* (measured between the C1 atoms of the sugar groups in U39 and G73 and more closely correlated to the switching sequence configuration), in each buffer condition (apo, Mg^2+^, TPP, and Mg^2+^&TPP, Figure 6B).

The riboswitch’s aptamer domain was mostly open when simulated in the apo buffer, with a major basin centered at *D_inter-arm_* ~87.8 Å, and its switching sequence exhibited both closed-like and open-like configurations (Figure 6B). The major basin was found at *D_co-stack_* ~8 Å (labeled *β*, aptamer domain in open state with P1 switch helix ^co-^stacking at the junction of the expression platform, Figure 6B) and had a shoulder corresponding to the loss of the P1 switch helix co-stacking interaction with *D_co-stack_* ~17.1 Å (labeled *α*, aptamer domain in open state without P1 switch helix co-stacking, Figure 6B). Additionally, there was a weakly populated basin at *D_inter-arm_* ~54.0 Å and *D_co-stack_* ~8 Å (labeled *γ*, aptamer domain in a partially closed-like ensemble with co-stacking of the P1 switch helix, Figure 6B). Representative snapshots corresponding to the three intermediate ensembles are shown in Figure 7.

**Figure 7.**
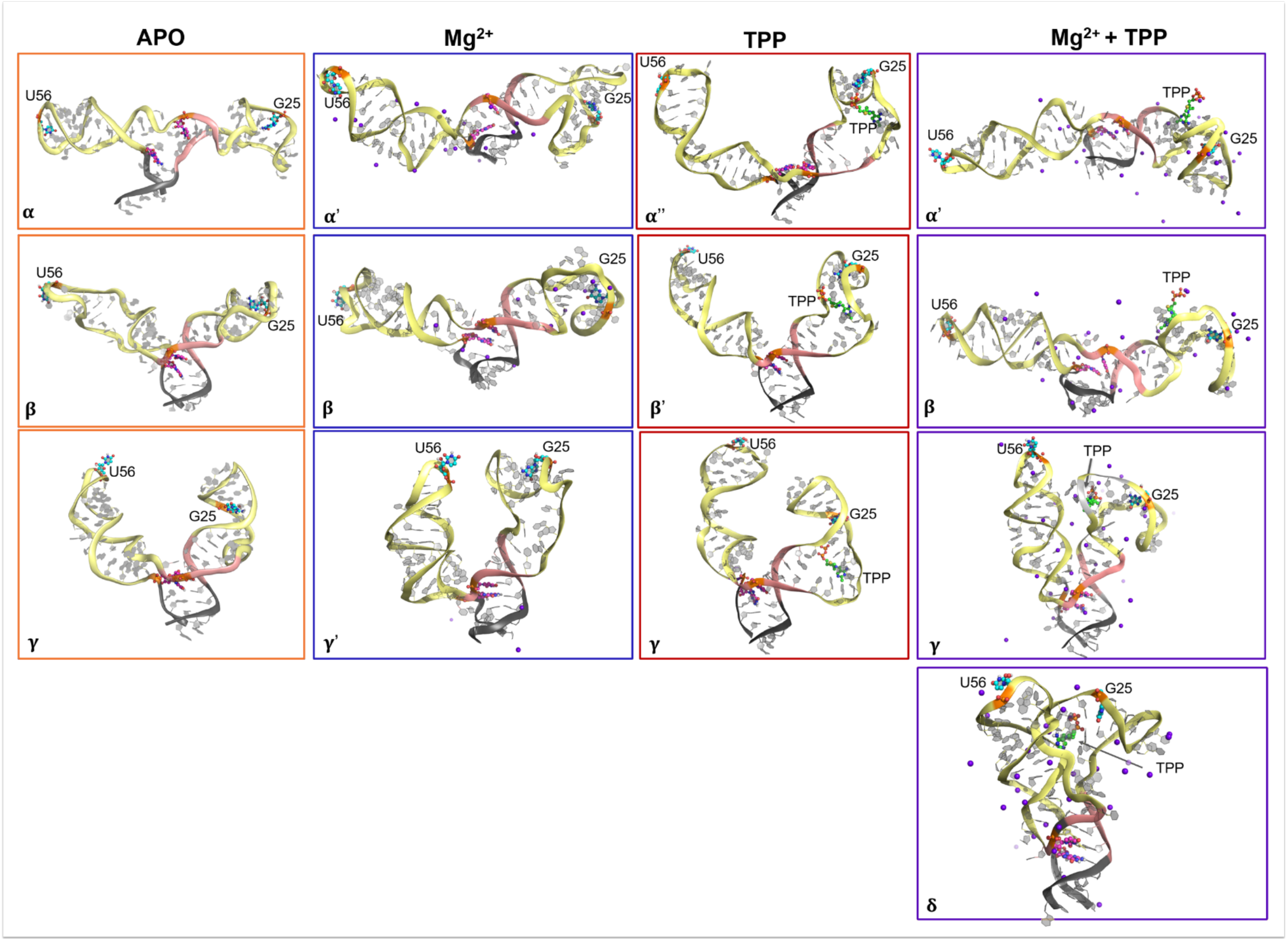
Representative snapshots of the riboswitch in APO (α, β, γ), Mg^2+^ (α’, β, γ’), TPP (β’, γ), and Mg^2+^&TPP (α’, β, δ) conditions. The states correspond to labeled basins in Figure. 6B. RNA was shown in cartoon representation, Mg^2+^ ions as purple spheres, and nucleotide pairs (G25, U56) and (U39, G73) highlighted in the ball-and-stick representation.

Simulations in the Mg^2+^ buffer showed three basins in the riboswitch conformational ensembles: the predominant basins *α*’ and *β* separated by a weak energy barrier and an isolated *γ’* basin. Compared to the *α* basin in the apo condition, *α*’ had a smaller co-stacking distance *D_co-stack_* ~10.9 Å in the Mg^2+^ buffer, and compared to the *γ* basin in the apo condition, *γ’* had a smaller inter-arm distance *D_inter-arm_* ~28.7 Å in the Mg^2+^ buffer (Figure 6B and snapshots in Figure 7).

Compared to the apo and Mg^2+^ buffers, molecular dynamic simulations of riboswitch with TPP alone resulted in a broader and shorter unimodal inter-arm distance distribution centered around ~69.4 Å and a slightly shorter co-stacking distance of ~7.0 Å *(β’*, Figure 6B, snapshots in Figure 7), but a very weakly populated, partially closed aptamer domain state strongly resembling the *γ* ensemble in the apo buffer. We also observed a weakly populated ensemble like the open aptamer domain state without co-stacking of the switching sequence (*α* basin, Figure 6B, snapshots in Figure 7).

In the Mg^2+^ & TPP buffer simulations, the riboswitch was mostly in a closed-like ensemble with inter-arm distance ~26.3 Å and co-stacking distance ~6.7 Å (*δ*-basin, Figure 6B and corresponding conformation in Figure 7), but also showed weakly populated ensembles corresponding to *α’, β*, and *γ* basins. Hence, both TPP and Mg^2+^ shifted RNA conformations toward compact states with shorter inter-arm and co-stacking distances, but the extent of this compaction and resulting intermediate ensembles were distinct. These findings are consistent with the FRET observations in Figure 4.

### Sampling configurational ensemble in unfolding pathway

To confirm that the rapid dynamics observed with smFRET do not correspond to unfolding transitions of RNA helices, we proceeded to measure the unfolding of the aptamer domain of the TPP riboswitch under force (Methods) in the same buffer conditions as in the smFRET experiments. Using a single-bead optical tweezer assay (Methods and SI Figure S7), we applied the force necessary to fully unfold the TPP riboswitch. We considered the TPP riboswitch to be fully unfolded once the full extension of the TPP riboswitch is reached, at which point the stretching behavior matches that of a worm-like chain polymer and the complete loss of secondary structure is assured (Figure 8A). We observed that the fraction of molecules that fully unfolded depended on the buffer conditions. In buffers containing TPP (TPP and TPP & Mg^2+^ buffers, Methods), the TPP riboswitch showed significantly more full unfolding events than under Apo or Mg^2+^-only conditions (Figure 8B). These results are consistent with the results obtained by smFRET in the apo and Mg^2+^ conditions, in which approximately 70% of the population showed no- FRET and approximately 30% exhibited FRET (Figure 4A), and in the TPP and Mg^2+^ & TPP buffers, in which the population that exhibited FRET was significantly higher than in the apo and Mg^2+^ buffers (Figure 4A). These results suggest that the states observed in the FRET-exhibiting populations (states *a*-*d* in Figure 4) do correspond to dynamic aptamer domain configuration states in which the riboswitch is predominantly folded (as in Figure 1).

**Figure 8.**
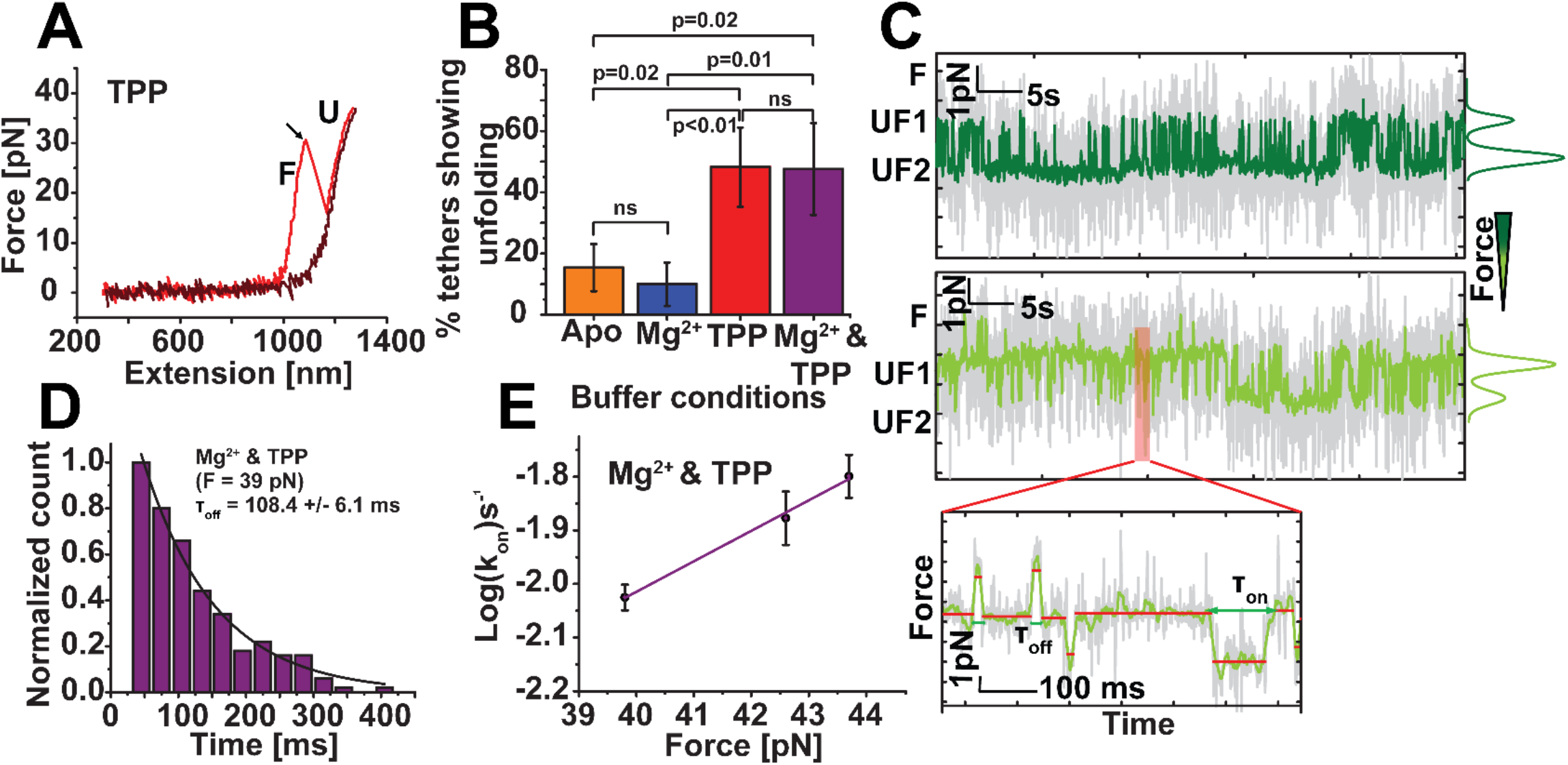
TPP riboswitch folding and unfolding kinetics. (A) A typical force extension curve (FEC) of the TPP riboswitch construct, in this case in the presence of 0.5 mM TPP. The arrow represents an unfolding event, as characterized by a sudden drop in the force corresponding with an increase in the length of the tether. The red FEC represents stretching of a construct with a folded TPP riboswitch, and the black one represents stretching of the construct with an unfolded TPP riboswitch. (B) Fraction of the stretched TPP riboswitch constructs showing unfolding events without ligand (apo, N = 26), with 1 M Mg^2+^ (N = 20), with 0.5 mM TPP (N = 27), and with Mg^2+^ + TPP (N = 21). The occurrence of unfolding events, i.e., the likelihood of finding folded TPP riboswitches, is significantly higher in the presence of TPP and TPP& Mg^2+^ as compared to either the presence of Mg^2+^ alone or the absence of ligand (apo). The error bars represent the relative standard deviation of counting statistics. (C) Example time-series force data showing rapid transitions between the folded and multiple unfolded states in the presence of 0.5 mM TPP, as determined by Gaussian mixture model analysis. Data was acquired using optical tweezer at 1 kHz (gray) and smoothed using a 25 ms moving average filter (green). Sudden increases and decreases in force identify the transitions between the folded (F) and various unfolded states (UF1 and UF2). The relative population of states is shown with Gaussian fits (to the right of time series data) for higher force (dark green, 20.0 pN) and lower force (light green, 18.6 pN). High-resolution time series trace (inset) obtained at approximately 18.6 pN pulling force. Folded states as fast as 25 ms and relatively long-lived unfolded states were detected at this force using change-point analysis. (D) Histogram of riboswitch folded state dwell time measured in the presence of 4.8 mM TPP and 0.5 M Mg^2+^ and at 39.8 pN force, as determined by change point analysis. The characteristic dwell time was τ_off_ = 108.4 ± 6.1 ms (single exponential decay fit constant ± std. error of the fit, N = 203). Histograms at additional forces found in SI Figure S8. (E) The plot of log_10_(k_on_) as a function of the force in the presence of 4.8 mM TPP and 0.5 M Mg^2+^ where k_on_ is inverse of τ_off_ (dwell time in the folded state, error bars represent the standard error of the exponential fits). Each data point (circle) represents 100 – 200 folded to unfolded transitions. The data were fitted using a weighted linear regression (solid line), and the dwell time at zero force was calculated to be τ_off_ = 17.2 ± 2.6 sec (y-intercept ± std. error of the fit).

To further study the unfolding transition, we held the TPP riboswitch in a passive optical trap with force applied. In this stressed condition, the riboswitch stochastically transitioned between multiple structural configurations, most likely representing transitions in P2/P4 stacking, which is correlated to P1 switching sequence base pairing and the extension of the 5′ end to the 3′ end of the riboswitch. In the example time-traces of TPP riboswitch switching sequence dynamics in TPP buffer held at 20.0 pN (Figure 8C, top) and 18.6 pN (Figure 8C, bottom), the TPP riboswitch extension exhibited fluctuations between multiple conformational states (also Figure 8C inset for detail). Lower forces shift the proportion of populated states to be more folded (Figure 8C, right for Gaussian fits of the histogram of state data). We used these time-trace data to determine the characteristic time of transitions between these states by fitting single exponential functions to dwell-time histograms acquired at various applied forces. For example, we found that the characteristic time of unfolding for the transition from the folded state to U1 (Figure 8C for states) was τ_off_(*F*) = 108.4 ± 6.1 ms at a tensile load of 39 pN for the TPP riboswitch in the Mg^2+^ & TPP buffer (Figure 8D). To determine the unfolding rate at zero force (see Methods for details), we measured these characteristic unfolding times at multiple forces (e.g. Figure 8C), calculated the corresponding characteristic times of unfolding (e.g. Figure 8D), generated a plot of log_10_(*k*_on_) as a function of force and extrapolated to zero force using Bell’s equation (Figure 8E). We found that the characteristic unfolding time at zero force was τ_off_ = 17.2 ± 2.6 s in the TPP & Mg^2+^ buffer (Figure 8E).

In total, our smFRET, DMD simulation, and optical tweezer results are consistent with a model of rapid ensemble switching underlying and tuning the transition between ON-like and OFF-like configuration ensembles, rather than a model that relies on the direct coupling between aptamer domain structural configuration state and switching sequence folding state that is long lived and binary.

## DISCUSSION

Our results suggest that the aptamer domain of the TPP riboswitch is a dynamic structure that is tuned by the presence of its co-ligands. Specifically, we found that the aptamer domain: *i*) exhibits dynamic processes occurring at sub-millisecond temporal scales; *ii*) is more likely to sample closed states in the presence of TPP, Mg^2+^, or both, *iii*) differentially responds to Mg^2+^ and to TPP, suggesting multiple closing pathways; and *iv*) requires non physiological conditions of Mg^2+^ in addition to TPP to generate a long lived OFF state, indicated by splitting of the ⟨τ_*D*(*A*)_⟩_*f*_ populations in the MFD plots (Figures 2–7). Further, aptamer domain dynamics occurs even in the presence of saturating amounts of TPP and Mg^2+^ that most favor the folded, OFF-like configurations (Figures 6–8), likely in part to Mg^2+^ allowing the riboswitch to sample intermediate ensembles (state *b*).

We further identified five conformational ensembles of the *Arabidopsis thaliana* TPP riboswitch’s aptamer domain as a function of Mg^2+^ and TPP ligand condition (*a, b, c, d*, and No-FRET, Figure 4). Each ligand condition gave rise to a unique subset of two aptamer domain conformational ensembles (Figure 4B) in addition to a significant population exhibiting open, no-FRET (Figure 4A) and unfolded (Figure 8B) configurations, which became smaller in the presence of ligands (Figures 4A and 8B). In the absence of ligand, the FRET-exhibiting aptamer domain population was predominantly in an open (*a*) configuration with a smaller fraction of the population in an intermediate (*c*) ensemble (Figure 4B). The presence of excess Mg^2+^ alone increased the relative population of the dynamic intermediate *c* ensemble but maintained a smaller population of the open *a* configuration (Figure 4B). A saturating concentration of TPP alone reduced aptamer sampling of the open *a* ensemble. Instead, it allowed the aptamer domain to sample the closed *d* state, sometimes exchanging with a second intermediate (*b*) ensemble (Figure 4B). In the presence of both ligands, the aptamer domain sampled both the *a* and *d* configurations, with the kinetics significantly favoring the closed *d* state (Figure 4B), thus suggesting that TPP introduces the OFF *(d)* state but that Mg^2+^ is required to stabilize it, due to the splitting of the ⟨τ_*D*(*A*)_⟩_*f*_ distribution in saturating Mg^2+^ & TPP conditions (Figure 3). Our computational modeling independently corroborates this ensemble switching scheme by identifying structural ensembles through clustering analysis (Figure 7) that correlate with the experimentally determined scheme (*α, β, γ*, and δ basins in the free energy landscape, respectively, Figure 6). Our results lead us to propose a new model for the conformational landscape of the TPP riboswitch’s aptamer domain as it binds to its Mg^2+^ and TPP ligands (Figure 9). Moreover, our results suggest the aptamer domain can take one of two independent closing pathways, depending on the order of ligand binding (Figure 9).

**Figure 9.**
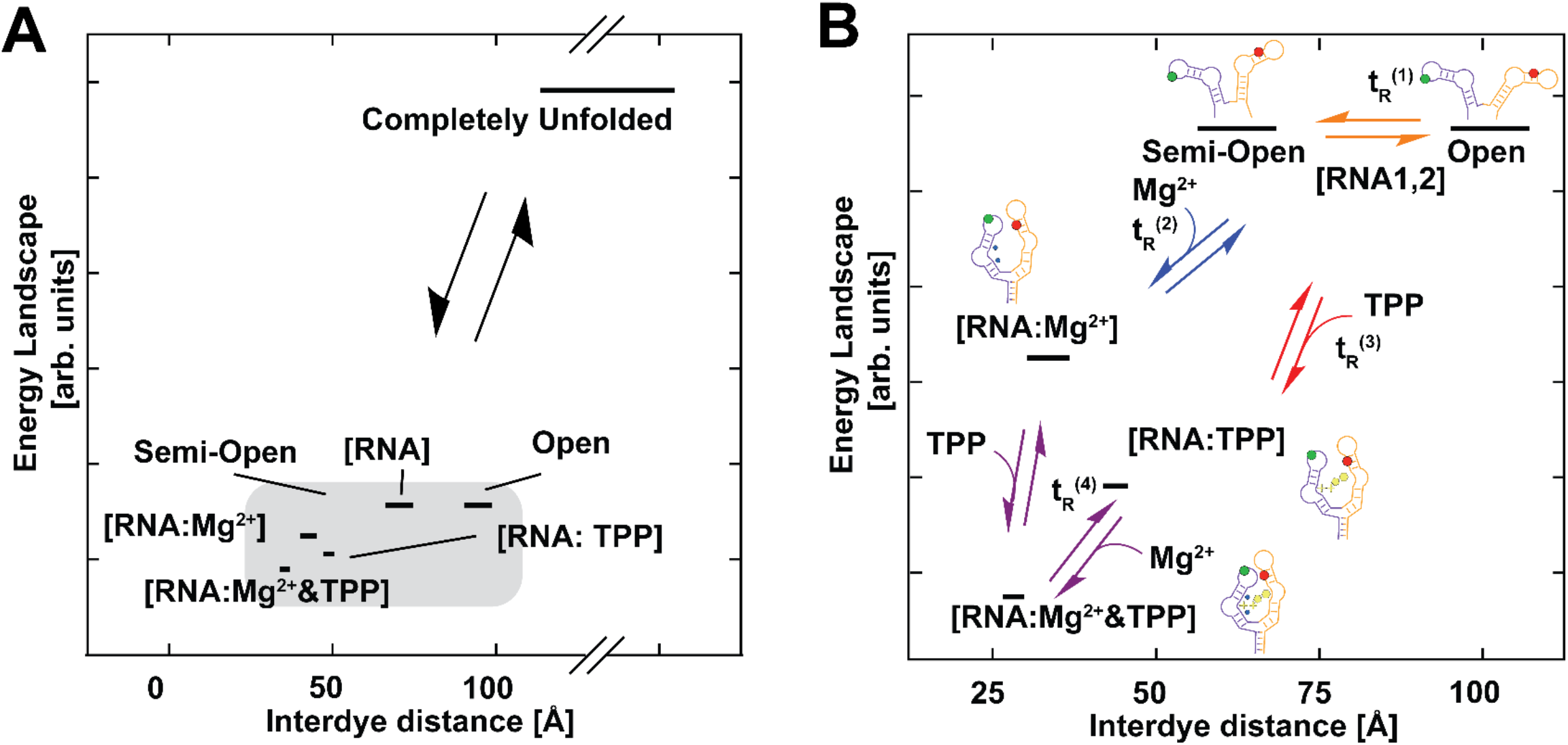
Energy Landscape of TPP Riboswitch Dynamics. Schematic of RNA closing. (A) The completely unfolded state accessible by tweezer experiments is shown along with the conformational states observed by smFRET. The long timescale of unfolding relative to kinetics observed by smFRET indicates that a much larger energy barrier must be overcome in these transitions. States observed by smFRET are highlighted in gray. (B) Zoomed representation of states highlighted in A in the context of Mg^2+^&TPP buffers. Two pathways exist from the open/semi-open state to the closed state. Conformational states present along the closing pathway of the riboswitch upon recognition of Mg^2+^ or TPP followed by completion of binding with both Mg^2+^&TPP. Labels for the times associated with each of these transitions are assigned based on fFCS.

RNA1 and RNA2 (Figure 9) ensembles share two exit pathways that go from open-like to closed-like configurational ensembles. RNA1 and RNA2 ensembles are distinct, extended (very low- or No-FRET) and share similar energies; thus, exchanging rapidly. RNA:Mg^2+^ corresponds to an intermediate configuration with fast dynamic motions due to the apparent weak effect of Mg^2+^ on distributing the configurational ensemble. RNA:TPP corresponds to another intermediate ensemble having undergone larger conformational changes. In the apo buffer conditions, the No-FRET population is largest (Figure 4A), and the lowest proportion of unfolding events are detected by optical tweezer experiments (Figure 8B), suggesting a relatively extended configuration. In these conditions, relaxation time *t*_R_^(1)^ in the μs time range (Figure 5C) and the long distance observed by FRET suggest that this open-like state is an ensemble of many possible unfolded configurations separated by low energy barriers, represented as RNA1 and RNA2 in Figure 9. The distinction between these ensembles is made to emphasize that TCSPC identified both a difficult to quantify No-FRET state and a low-FRET state. Relaxation time *t*_R_^(2)^ in the 10s of μs likely corresponds to the process of a Mg^2+^ ion binding and the corresponding local configurational changes, which leads to a more highly populated intermediate, semi-closed ensemble of the aptamer domain and a decreased No-FRET population as compared to apo buffer (Figure 4). Relaxation time *t*_R_^(3)^ is slower than *t*_R_^(2)^ indicating a larger energy barrier likely corresponding to transitions involving global conformational reorganization, like those associated with TPP binding that favors compact configurations (Figure 4). *t*_R_^(4)^ is the slowest relaxation time, likely corresponding to the limiting reaction step, transitions between the closed and intermediate configuration ensembles. This interpretation is corroborated by smMFD data (Figure 3) in saturating conditions of Mg^2+^ & TPP, in which the low-⟨τ_*D*(*A*)_⟩_*f*_, high-FRET population is separated from the other dynamic population. This type of separation indicates that exchange between these ensembles occurs on timescales comparable to or longer than the diffusion time, like *t*_R_^(4)^ The trend of faster correlation times, as well as changes in population fractions from TCSPC, in Mg^2+^, TPP, and Mg^2+^ & TPP buffers suggests that transitions toward the OFF-like and intermediate ensembles is only favored in these extreme conditions compared to the apo buffer or at low Mg^2+^ when both favor extended conformations. The fastest correlation times, exhibited in Mg^2+^ & TPP conditions, further suggest cooperativity between Mg^2+^ and TPP that favors transitions toward the long lived OFF state and allows sampling of both closing pathways and the corresponding configurations.

In addition to the ensemble switching in the aptamer domain, we found significantly slower dynamics in the switching sequence of helix P1. Under a relatively high (10s of pN) force, we observed dynamic switching between a fully- folded state (F), which likely corresponds to base pairing in the P1 helix to achieve the switching sequence OFF state, and two less-folded states (UF1, UF2), which likely correspond to the transition to the ON state by opening of the P1 helix (Figure 8C). Using Bell’s equation, we calculated that the equilibrium (zero-force) transition rate of the switching sequence between the OFF and ON state in the presence of both TPP and Mg^2+^ (Figure 8E) was 4 orders of magnitude slower than the slowest rate between structural configurations in the aptamer domain also in the presence of both TPP and Mg^2+^ (Figure 5C). This disparity in timescale indicates that, while TPP and Mg^2+^ recognition and binding in the aptamer domain certainly stabilize the P1 helix (Figures 6–8), transitions between the structural configuration ensembles in the aptamer domain cannot correspond one-to-one to unfolding, TPP riboswitch switching sequence ON/OFF transitions (Figure 9B). Thus, while the larger-scale ON/OFF behavior of the riboswitch takes place on longer timescales, fast ensemble switching in the aptamer domain may allow more subtle biasing of the riboswitch toward ON-like or OFF-like states with a rheostat-like tunability given by the switching kinetics.

The folding dynamics of aptamer domains are key to riboswitch function, and divalent ion co-ligand binding to the aptamer domain can facilitate this folding (28). Further, the TPP riboswitch recognizes TPP in a step-wise fashion mediated by Mg^2+^ (27). Our results suggest that only through coordinating with a divalent ion, Mg^2+^, is TPP able to stabilize a closed aptamer domain’s tertiary structure (Figures 3, 6, and 7). Further, all four observed, distinct timescales occur in the sub-millisecond range, which is well below the timescales associated with slow, high-energy conformational changes or transcriptional folding kinetics (Figures 5, 6, and 8). Thus, the short-timescale dynamics of Mg^2+^-aptamer domain binding regulate the longer-timescale switching sequence ON/OFF conformational transitions, giving rise to a stepwise closing pathway. Our model also resolves an apparent disparity between the previously suggested fast or slow binding recognition mechanisms (17,19), where the fast kinetics explains the ensemble switching between all the aptamer domain configuration that may correlate with, but need not induce, switching sequence and expression domain state transitions that occur at slower timescales. Because changes in Mg^2+^ and TPP concentrations regulate fast structural state kinetics to favor subsets of configurational ensembles, which do not correlate one-to-one with the ON/OFF states, we hypothesize a model where the TPP riboswitch function can be finely-tuned via kinetics rheostat mechanism rather than a binary open/closed aptamer domain configurations that directly correlating to the ON/OFF regulation of gene expression. We suggest that such rheostatic tuning could allow for fast response times for regulation of TPP concentrations in cells.

## ACKNOWLEDGEMENTS

We thank Thomas O. Peulen for developing ChiSurf and making it available for data analysis at https://github.com/Fluorescence-Tools. smFRET software analysis can be found at http://www.mpc.hhu.de/software/software-package.html. This work was supported by NSF (CAREER MCB 1749778 to HS, CBET-1553945 to FD), NIH (SC TRIMH COBRE 1P20GM130451, 2R01MH081923-11A1, EPIC COBRE P20GM109094, 1R15AI137979, and R35GM119691), start-up funds from Clemson University (HS, JA), Creative Inquiry at Clemson University, and the Center for Optical Materials Science and Engineering Technologies (COMSET). The authors declare that they have no conflicts of interest regarding the contents of this article. The content is solely the responsibility of the authors.

## AUTHOR CONTRIBUTIONS

FD, JA, and HS conceived and designed the project. JM and GH measured and analyzed smFRET experiments. NS and FD performed DMD simulations. SG and JA performed and analyzed the optical tweezer experiments. JM, GH, FD, JA, SG, and HS wrote the manuscript.

## DATA AVAILABILITY

The authors declare that all data supporting the findings of this study are available within the paper and its SI.

