## Supplemental Information for "Ensemble Switching Unveils a Kinetic Rheostat Mechanism of the Eukaryotic Thiamine Pyrophosphate Riboswitch"

Junyan Ma[1,2,*§], Nabanita Saikia[3,*], Subash Godar[3,*], George L. Hamilton[3], Feng Ding[3], Joshua Alper[3,4,5], Hugo Sanabria[2,3]
[1]Department of Chemistry, Clemson University, Clemson, S.C., USA
[2]Center for Optical Materials Science and Engineering Technologies, Clemson, S.C., USA. [3]Department of Physics and Astronomy, Clemson University, Clemson, S.C., USA [4]Department of Biological Sciences, Clemson University, Clemson, S.C., USA
[5]Eukaryotic Pathogen Innovation Center, Clemson University, Clemson, S.C., USA

To whom correspondence should be addressed: Hugo Sanabria, Clemson University,, Joshua Alper, Clemson University,, Feng Ding, Clemson University,
* These authors contributed equally
§ Current affiliation: College of Chemistry and Chemical Engineering, Anyang Normal University, Anyang, P. R. China.

### SUPPLEMENTAL INFORMATION

### SUPPLEMENTARY NOTES

#### Acceptor quenching in the presence of $Mg^{2+}$ and TPP

Static quenching refers to the process by which a fluorophore is rendered non-fluorescent due to the formation of a complex with another molecule, inducing a new, stable ground state. Decays to this ground state are non-photon-emitting. This is often due to hydrophobically-induced stacking with elements of the labeled macromolecule near the labeling site. This form of quenching does not affect the fluorescence lifetime, however, as those fluorophores do not emit any photons while unaffected fluorophores emit normally. However, this results in a net loss of fluorescence signal; thus, the mean of $F_D/F_A$ is affected if either donor or fluorophore is more likely to experience static quenching. Thus, changes in FRET efficiency are tractable most easily through the change in the donor fluorescence lifetime. Static quenching differs from the case of dynamic quenching which results from processes such as collisions of the fluorophore with "quencher" molecules. Dynamic quenching refers quenching by additional, non-radiative decay pathways that effectively increase the decay rate of the excited-state population (thus reducing the fluorescence lifetime). In our case, we identified that in the closed configuration (High-FRET), there is significant acceptor quenching that shifts the population upward over the expected $F_D/F_A$ ratio as given by the static FRET line. For that reason, we used two distinct FRET lines (Figure 2D). The corrected FRET line for the acceptor quenching overlays the High-FRET population. Note that the $F_D/F_A$ ratio reports on the fluorescence of the donor and acceptor, but $\langle\tau_{D(A)}\rangle_f$ only reports on the donor quenching by FRET, without considering the emission of the acceptor. This is one of the reasons why MFS is uniquely poised to identify and quantify potential photophysical problems. For this reason, in this study we focus our analysis on the one-dimensional projection of the $\langle\tau_{D(A)}\rangle_f$.

### SUPPLEMENTARY METHODS

#### Sequences for Optical Tweezer DNA Extensions

The RNA-flanking DNA extension sequence is
5’ GGTATTAACGCCGCCTCACTTTTGTGGCGTTAGGTauauau TPP RIBOSWITCH uaacaa CACGACGCCGATGGGTACCGCATCCCCCTTTCGCCA 3’

5' GGTATTAACGCCGCCTCACTTTTGTGGCGTTAGGT auauau RNA uaacaa CACGACGCCGATGGGTACCGCATCCCCCTTTCGCCA 3'

where capital letter nucleotides represent DNA bases. For each handle, the primer sequences are given by

*Handle* 1 – Forward primer: 5' N CGCCGATCAACTGGGTGCCA 3'
*Handle* 1 – Reverse primer: 5' ACCTAACGCCACAAAAGTGAGGCGGCGTTAATACC F TGCGCT 3'
*Handle*2-Forward primer: 5' CACGACGCCGATGGGTACCGCATCCCCCTTTCGCCAG*C*T*GG 3'
*Handle* 2 - Reverse primer: 5' b CGGCGGGTGTGGTGGTTACG 3'

where N refers to amine group modification, F refers to a tetrahydrofuran abasic site, * refers to a phosphorothioate bond modification, and b refers to a biotin modification.

**FRET with Multiparameter Fluorescence Spectroscopy**

$F_{D|D}$ and $F_{A|D}$ are the signals to the donor and acceptor detection channels (green and red, respectively) following donor excitation, corrected for their corresponding detection efficiencies and backgrounds. The ratio of donor over acceptor fluorescence is related to FRET efficiency as $\frac{F_{D|D}}{F_{A|D}} = \frac{\Phi_{FA}}{\Phi_{FD(0)}}(E^{-1} - 1)$, where $\Phi_{FD(0)}$ and $\Phi_{FA}$are the donor and acceptor quantum yields. Therefore, when $\frac{F_{D|D}}{F_{A|D}} \rightarrow \infty$ , then the FRET efficiency, $E$, approaches 0, and when $\frac{F_{D|D}}{F_{A|D}} \rightarrow 0$, then $E \sim 1$. $\langle\tau_{D(A)}\rangle_f$ is the burst-integrated fluorescence lifetime of the donor fluorophore, as determined by maximum likelihood estimation for each burst. $E$ is related to $\langle\tau_{D(A)}\rangle_f$ by $E = 1 - \langle\tau_{D(A)}\rangle_f / \langle\tau_{D(0)}\rangle_f$ in the case of static FRET states.

Obtaining information about dynamic processes at sub-millisecond timescales is challenging, even for smFRET experiments (1). To overcome these challenges, we use MFS and its ability to map biomolecular dynamics effects over several orders of magnitude in time (2-7). For example, if molecules sample multiple conformational states during the observation time, a single average distribution over the FRET intensity indicators $F_D/F_A$ and the $\langle\tau_{D(A)}\rangle_f$ will appear. On the contrary, if conformational states are stable for times longer than the observation time (burst duration), isolated populations corresponding to each state will show up. Moreover, due to changes in the first and second moments of the fluorescence lifetime distribution in the dynamic case that the FRET intensity indicators $F_D/F_A$ and $\langle\tau_{D(A)}\rangle_f$ will differentially reflect, the weighted mean of the donor lifetime for exchanging states is biased toward longer lifetimes. Thus, in multidimensional histograms, the observed population deviates from the expected relationship for static states that follow Förster theory, called the static FRET line. Therefore, plotting $F_D/F_A$ against the average fluorescence lifetime $\langle\tau_{D(A)}\rangle_f$ per single-molecule event, as done in Figure 2, serves as the first visual indicator of conformational dynamics.

Our implementation of MFS with pulsed-interleaved excitation (PIE) also allows the use of apparent stoichiometry, $S_{PIE}$, to quantify the donor-to-acceptor stoichiometry for each molecule. $S_{PIE}$ is the ratio of photon counts following donor excitation to the total number of photon counts after both excitation pulses, normalized according to the fluorophore quantum yields and the ratio of donor to acceptor excitation intensities and corrected for background, crosstalk from the donor to acceptor channel, and direct excitation of the acceptor by the donor excitation laser (8).

$$S_{PIE} = \frac{F_{A|D} + F_{D|D}}{F_{A|D} + F_{D|D} + F_{A|A}} \tag{1}$$

**Ensemble Time-Correlated Single Photon Counting (eTCSPC) analysis**

Time-resolved fluorescence decays ($F(t)$) were modeled using a multi-exponential model given by

$$F(t) = \sum_i x^{(i)} \exp(-t/\tau^{(i)}) \tag{2}$$

where $x^{(i)}$ is the corresponding population fraction, and $\tau^{(i)}$ is the fluorescence lifetime. Here, we used a two-exponential model to fit TCSPC data, resulting in a reasonably low $\chi^2$. The averaged species lifetimes and average fluorescence lifetimes are

$$\langle\tau\rangle_x = \sum_i x^{(i)} \cdot \tau^{(i)} \text{ , and}$$
$$\langle\tau\rangle_f = \frac{\sum_i x^{(i)} \cdot \left(\tau^{(i)}\right)^2}{\langle\tau\rangle_x}, \quad (3)$$

respectively.

To model a superposition of $j$ Gaussian-distributed FRET states, with population fractions $x_{DA}^{(j)}$, one needs to generate an interdye distance distribution $p(R_{DA})$ following

$$p(R_{DA}) = \sum_j \; x_{DA}^{(j)} \frac{1}{\sqrt{2\pi} \cdot \sigma_{DA}} \exp\left(-\frac{\left(R_{DA} - \langle R_{DA}^{(j)}\rangle\right)^2}{2\,\sigma_{DA}^2}\right), \quad (4)$$

where $\sigma_{DA}$ is the width of the distribution determined by the Accessible Volume of the fluorescent dyes. $\langle R_{DA}\rangle$ is the mean interdye distance.

*Filtered Fluorescence Correlation Spectroscopy*

To properly fit the species auto- and cross-correlation function we used a set of equations given by

$$G_{i,i}(t_c) = 1 + \frac{1}{N_{Br}} \cdot G_{diff}^{(i)}(t_c) \cdot G_T^{(i)}(t_c) \cdot G_B^{(i)}(t_c) \cdot \left[1 + \sum_{y=1}^{5} AC_{i,i}^{(R)} \;(\exp\left(-\frac{t_c}{t_R^{(y)}}\right) - 1)\right]$$
$$G_{m,m}(t_c) = 1 + \frac{1}{N_{Br}} \cdot G_{diff}^{(m)}(t_c) \cdot G_T^{(m)}(t_c) \cdot G_B^{(m)}(t_c) \cdot \left[1 + \sum_{y=1}^{4} AC_{m,m}^{(R)} \;(\exp\left(-\frac{t_c}{t_R^{(y)}}\right) - 1)\right] \quad (5)$$
$$G_{i,m}(t_c) = 1 + \frac{1}{N_{Br}} \cdot G_{diff}^{(i,m)}(t_c) \cdot G_B^{(x)}(t_c) \cdot \left[1 - CA_{i,m} \sum_{y=1}^{4} CC_{i,m}^{(R)} \;\exp\left(-\frac{t_c}{t_R^{(y)}}\right)\right]$$

where $t_R^{(y)}$ are the relaxation times that correspond to the exchange times between selected filtered species ($y$) with corresponding absolute amplitudes of the species auto-correlation functions (sACF) $AC_{i,i}^{(R)}$and the relative normalized amplitudes of the species cross-correlation function (sCCF) $CC_{i,m}^{(R)}$, normalized to the absolute amplitude $CA_{i,m}$. The contribution to the correlation function from the triplet state or dark state kinetics of the dyes is $G_T^{(x)}(t_c) = \left[1 - T^{(x)} + T^{(x)} \cdot \exp\left(-\frac{t_c}{t_T^{(x)}}\right)\right]$, where $T^{(i)}$ is the triplet state dynamics amplitude. $N_{Br}$ is the average number of bright molecules in the focus corresponding to the sACF, and $N_{CC}$ in the sCCF corresponds to the inverse of the initial amplitude $G_{i,m}^{(x)}(0)$. $G_B^{(i)}(t_c)$ is defined for the bleaching term as $G_B^{(i)}(t_c) = \left[1 - B^{(i)} + B^{(i)} \cdot \exp\left(-\frac{t_c}{t_B^{(i)}}\right)\right]$, where $B$ is a fractional contribution to the correlation amplitude.

$G_{diff}^{(i)}(t_c)$ is the diffusion term of species $x$:

$$G_{diff}^{(i)}(t_c) = \left(1 + \frac{t_c}{t_{diff}^{(i)}}\right)^{-1} \cdot \left(1 + \left(\frac{\omega_0}{z_0}\right)^2 \cdot \left(\frac{t_c}{t_{diff}^{(i)}}\right)\right)^{-1/2}, \quad (6)$$

where $\frac{\omega_0}{z_0}$ is a geometric parameter defining the confocal illumination volume, and $t_{diff}^{(i)}$ is the characteristic time of diffusion. To model the conformational switching, we needed four relaxation times for both the sACF and sCCF. This model complements the limiting states observed in TCSPC experiments.

#### Error propagation

To estimate the statistical uncertainty for the determined distances, we use an error propagation rule which considers the uncertainty of the optimization algorithm while minimizing the figure of merit, $\chi^2$ ($\Delta\chi^2$), and the uncertainty associated with the orientation of the dyes, $\kappa^2$ ($\Delta\kappa^2$). For $\Delta\chi^2$, we first calculate the maximum allowed $\chi^2_{r,\max}$ for a given confidence level (*P;* e.g., for $2\sigma$ or $P = 0.95$), given by

$$\chi^2_{r,\max}(P) = \chi^2_{r,\min} \cdot \left[1 + n/\upsilon \cdot cdf^{-1}(F(n, \upsilon, P))\right] \quad (7)$$

The minimum, $\chi^2_{r,\min}$, is obtained from the fit for each of the identified distances, $R_{DA}$. Here, $n$ is the number of data points fit, $\upsilon$ is the number of degrees of freedom in the fit, and $cdf^{-1}(F(n, \upsilon, P))$ is the inverse of the cumulative distribution for the F-distribution for given $n$, $\upsilon$, and $P$. It is worth noting that increasing the number of states in the model function tends to reduce $\chi^2$ through an increase in $\upsilon$, as each additional term introduces additional fit parameters. To avoid over-fitting in this way, we only accept the addition of a term corresponding to an additional state if it results in significant improvement to $\chi^2$ compared to the model with one fewer state. $\Delta\kappa^2$ is determined based on the arithmetic mean of the $\kappa^2$ distribution. For the overall uncertainty, we apply the following equation:

$$\varepsilon(\kappa^2, k_{FRET}) = \Delta R^2_{DA}(\kappa^2) + \Delta R^2_{DA}(k_{FRET}), \quad (8)$$

where $\Delta R^2_{DA}(\kappa^2)$ and $\Delta R^2_{DA}(k_{FRET})$ are the distance uncertainty contributions due to the orientation factor and the width of the FRET distance distribution, respectively.

### SUPPLEMENTARY TABLES AND FIGURES

**Table S1.** System setup with #counterions in APO, $Mg^{2+}$, TPP, and $Mg^{2+}$&TPP conditions for the riboswitch in REXDMD simulations.

| *# counterions* | *APO* | *$Mg^{2+}$* | *TPP* | *$Mg^{2+}$&TPP* |
|---|---|---|---|---|
| ***$Na^+$*** | 77 | 1 | 80 | 2 |
| ***$Mg^{2+}$*** | - | 38 | - | 39 |

**Table S2.** Static FRET Lines

A) Line equations

| Sample | FRET lines (x=⟨ $\tau_{D(A)}$⟩f) |
|---|---|
| Apo | (0.7552/0.4700)/((3.7760/((-0.0427*x^3)+(0.2999*x^2)+0.4862*x+-0.0415))-1) |
| Mg | (0.7220/0.3800)/((3.6100/((-0.0471*x^3)+(0.3170*x^2)+0.4805*x+-0.0402))-1) |
| TPP | (0.6105/0.3200)/((3.7576/((-0.0441*x^3)+(0.2327*x^2)+0.7660*x+-0.0712))-1) |
| TPP&$Mg^{2+}$ | (0.7300/0.4800)/((3.6500/((-0.0460*x^3)+(0.3127*x^2)+0.4819*x+-0.0405))-1)<br>(0.7296/0.0100)/((3.6480/((-0.0461*x^3)+(0.3129*x^2)+0.4818*x+-0.0405))-1) |

B) Correction Parameters

| Sample | $\Phi_D$ | $\Phi_A$ | $\tau_{D0}$ [ns] | Detection Eff. Ratio, G/R | Crosstalk G→R (%) | Prob. Of Acceptor Excitation by Donor Laser (%) | $BG_G$ [kHz] | $BG_R$ [kHz] |
|---|---|---|---|---|---|---|---|---|
| Apo | 0.7552 | 0.4700 | 3.7760 | 3.7 | .017 | .02 | .640 | .369 |
| Mg | 0.7220 | 0.3800 | 3.6100 | 3.7 | .017 | .02 | .651 | .381 |
| TPP | 0.7300 | 0.3200 | 3.6480 | 3.7 | .017 | .02 | .670 | .370 |
| TPP&$Mg^{2+}$ | 0.7300 | 0.4800 | 3.6480 | 3.7 | .017 | .02 | .435 | .305 |

**Table S3.** $\chi^2$improvement upon the addition of free parameters.

| Sample | One Gaussian $\chi^2$ | Two Gaussian $\chi^2$ | Three Gaussian $\chi^2$ |
|---|---|---|---|
| Apo | 6.8 | 2.5 | 2.1 |
| Mg | 15.0 | 2.4 | 2.1 |
| TPP | 3.6 | 2.0 | 2.0 |
| TPP&$Mg^{2+}$ | 9.3 | 2.8 | 2.6 |

**Table S4.** Ensemble Time-Correlated Single Photon Counting
A) Donor only Lifetime Decay

| **Species** | **$\tau_1$ (ns)** | **$x_1$** | **$\tau_2$ (ns)** | **$x_2$** | **$\tau_3$ (ns)** | **$x_3$** | **$\tau_4$ (ns)** | **$x_4$** | **$\langle\tau\rangle_x$ (ns)** | **$\langle\tau\rangle_f$ (ns)** | **$\chi_r^2$** |
|---|---|---|---|---|---|---|---|---|---|---|---|
| Apo | 3.95 | 0.59 | 2.42 | 0.14 | 0.79 | 0.11 | 0.09 | 0.16 | 2.771 | 3.618 | 2.01 |
| TPP | 3.85 | 0.54 | 2.29 | 0.15 | 0.80 | 0.12 | 0.09 | 0.19 | 2.536 | 3.492 | 2.23 |
| Mg | 4.18 | 0.53 | 2.69 | 0.17 | 0.82 | 0.11 | 0.10 | 0.19 | 2.781 | 3.822 | 2.75 |
| TPP&$Mg^{2+}$ | 4.04 | 0.57 | 2.77 | 0.17 | 0.88 | 0.09 | 0.10 | 0.17 | 2.867 | 3.724 | 2.21 |

B) Donor Acceptor Distances and fractions as model with a two Gaussian distributed state model

| **Species** | **$R_{DA1}$(Å) ± ε** | **$\sigma_1$** | **$x_1$** | **$R_{DA2}$(Å) ± ε** | **$\sigma_2$** | **$x_2$** | **$\chi_r^2$** | **Donly** |
|---|---|---|---|---|---|---|---|---|
| Apo | 33.90±2.3 | 6 | 0.26 | 66.42±4.5 | 6 | 0.74 | 2.52 | 0.73 |
| Mg | 31.71±1.9 | 6 | 0.61 | 58.67±3.3 | 6 | 0.39 | 2.40 | 0.67 |
| TPP | 25.36±1.0 | 6 | 0.78 | 43.26±1.2 | 6 | 0.22 | 2.00 | 0.59 |
| TPP&$Mg^{2+}$ | 26.08±1.2 | 6 | 0.83 | 55.63±2.5 | 6 | 0.17 | 2.83 | 0.59 |

C) *f*FCS fit parameters. τ in units ms.

| AC | | | | | | | | | | | | | | | | | | |
|---|---|---|---|---|---|---|---|---|---|---|---|---|---|---|---|---|---|---|
| Sample | Correlation | N | $(w_0/z_0)^2$ | $\tau_{diff}$ | T | $\tau_T$ | $B_i$ | $\tau_B$ | $AC_1$ | $\tau_{R1}$ | $AC_2$ | $\tau_{R2}$ | $AC_3$ | $\tau_{R3}$ | $AC_4$ | $\tau_{R4}$ | $AC_5$ | $\tau_{R5}$ |
| apo | LF-LF | 1.062 | 4.415 | 0.774 | 0.095 | 0.000 | 0.132 | 0.004 | 0.005 | 0.002 | 0.052 | 0.050 | 0.012 | 0.435 | 0.000 | 3.246 | 0.000 | 1.626 |
| | HF-HF | 1.062 | 4.415 | 0.774 | 0.095 | 0.000 | 0.132 | 0.004 | 0.005 | 0.002 | 0.052 | 0.050 | 0.012 | 0.435 | 0.000 | 3.246 | 0.000 | 1.626 |
| Mg2+ | LF-LF | 0.429 | 4.415 | 1.186 | 0.255 | 0.000 | 0.100 | 0.010 | 0.069 | 0.004 | 0.098 | 0.080 | 0.042 | 1.150 | 0.000 | 3.219 | 0.000 | 10.598 |
| | HF-HF | 1.051 | 4.415 | 1.186 | 0.131 | 0.000 | 0.109 | 0.010 | 0.107 | 0.004 | 0.050 | 0.080 | 0.036 | 1.150 | 0.000 | 3.219 | 0.000 | 10.598 |
| TPP | LF-LF | 0.528 | 4.415 | 0.671 | 0.000 | 0.003 | 0.000 | 0.050 | 0.278 | 0.001 | 0.188 | 0.008 | 0.099 | 0.098 | 0.000 | 2.381 | 0.000 | 11.033 |
| | HF-HF | 1.024 | 4.415 | 0.671 | 0.092 | 0.003 | 0.081 | 0.050 | 0.118 | 0.001 | 0.088 | 0.008 | 0.000 | 0.098 | 0.000 | 2.381 | 0.000 | 11.033 |
| Mg2+ & TPP | LF-LF | 0.325 | 4.415 | 1.165 | 0.191 | 0.000 | 0.011 | 0.006 | 0.167 | 0.000 | 0.221 | 0.003 | 0.109 | 0.072 | 0.018 | 1.521 | 0.000 | 1.883 |
| | HF-HF | 1.072 | 4.415 | 1.165 | 0.101 | 0.000 | 0.156 | 0.006 | 0.084 | 0.000 | 0.049 | 0.003 | 0.088 | 0.072 | 0.018 | 1.521 | 0.000 | 1.883 |
| CC | | | | | | | | | | | | | | | | | | |
| Sample | Correlation | N | $(w_0/z_0)^2$ | $\tau_{diff}$ | | | $B_i$ | $\tau_B$ | $CC_1$ | $\tau_{R1}$ | $CC_2$ | $\tau_{Rw}$ | $CC_3$ | $\tau_{R3}$ | $CC_4$ | $\tau_{R4}$ | CA | |
| apo | LF-HF | 3.491 | 4.415 | 0.774 | | | 0.336 | 1.626 | 0.194 | 0.002 | 0.089 | 0.050 | 0.203 | 0.435 | 0.515 | 3.246 | 1.357 | |
| | HF-LF | 2.220 | 4.415 | 0.774 | | | 0.000 | 1.626 | 0.194 | 0.002 | 0.089 | 0.050 | 0.203 | 0.435 | 0.515 | 3.246 | 1.356 | |
| Mg2+ | LF-HF | 3.443 | 4.415 | 1.186 | | | 0.314 | 3.219 | 0.235 | 0.004 | 0.089 | 0.080 | 0.219 | 1.150 | 0.463 | 10.598 | 1.186 | |
| | HF-LF | 2.401 | 4.415 | 1.186 | | | 0.000 | 3.219 | 0.235 | 0.004 | 0.089 | 0.080 | 0.219 | 1.150 | 0.463 | 10.598 | 1.134 | |
| TPP | LF-HF | 6.541 | 4.415 | 0.671 | | | 0.083 | 11.033 | 0.450 | 0.001 | 0.205 | 0.008 | 0.121 | 0.098 | 0.224 | 2.381 | 2.258 | |
| | HF-LF | 6.446 | 4.415 | 0.671 | | | 0.000 | 11.033 | 0.450 | 0.001 | 0.205 | 0.008 | 0.121 | 0.098 | 0.224 | 2.381 | 2.197 | |
| Mg2+ & TPP | LF-HF | 6.057 | 4.415 | 1.165 | | | 0.417 | 1.521 | 0.628 | 0.000 | 0.148 | 0.003 | 0.051 | 0.071 | 0.173 | 1.883 | 3.937 | |
| | HF-LF | 5.523 | 4.415 | 1.165 | | | 0.314 | 1.521 | 0.628 | 1.000 | 0.148 | 1.003 | 0.051 | 1.071 | 0.173 | 2.883 | 4.034 | |

D) The value of $\chi^2$ for each Gaussian distribution model.

| Sample | One Gaussian $\chi^2$ | Two Gaussian $\chi^2$ | Three Gaussian $\chi^2$ |
|---|---|---|---|
| Apo | 6.8 | 2.5 | 2.1 |
| Mg | 15.0 | 2.4 | 2.1 |
| TPP | 3.6 | 2.0 | 1.9 |
| TPP&$Mg^{2+}$ | 9.3 | 2.8 | 2.6 |

E) The diffusion time in each condition

| Species | $t_{diff}$ |
|---|---|
| Apo | 0.91 |
| Mg | 1.16 |
| TPP | 1.55 |
| TPP&$Mg^{2+}$ | 2.60 |

F) Anisotropy values of each sample

| Sample/Anisotropy | $r_{donly}^{(GG)}$ | $r_{DA}^{(GR)}$ | $r_{DA}^{(AA)}$ |
|---|---|---|---|
| Apo | 0.13 | 0.05 | 0.34 |
| Mg | 0.11 | 0.16 | 0.43 |
| TPP | 0.18 | 0.13 | 0.16 |
| TPP&$Mg^{2+}$ | 0.18 | 0.19 | 0.34 |

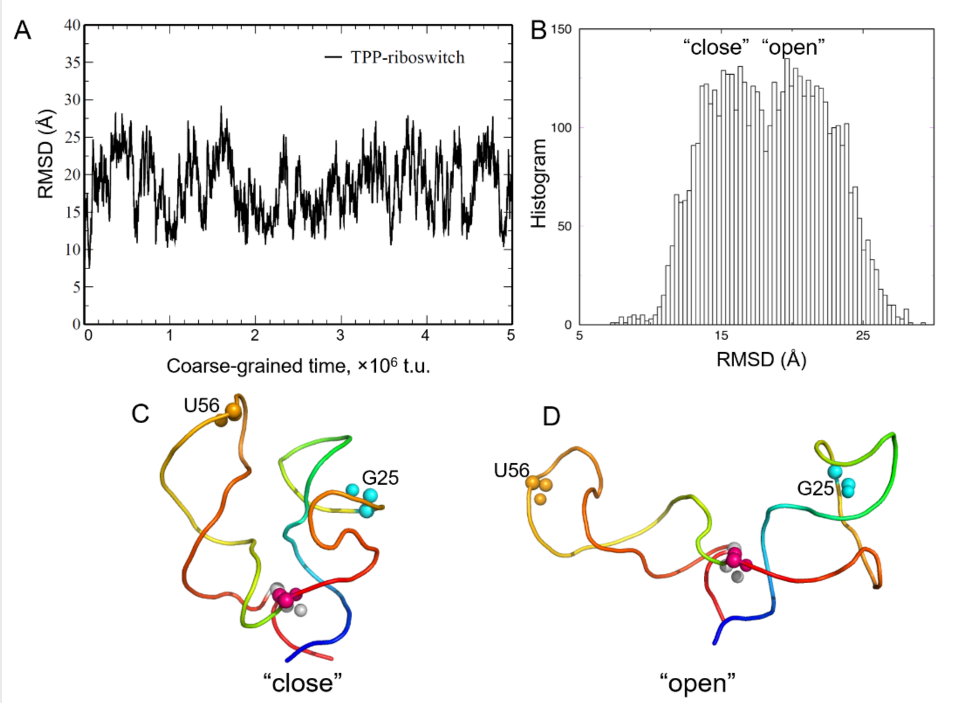


**Figure S1.** (A) RMSD as a function of coarse-grained time for TPP-riboswitch. (B) Histogram of RMSD depicting the "close" and "open" states for TPP-riboswitch. (C–D) Representative conformations of "close" and "open" states in TPP-riboswitch. The blue and red loop regions correspond to the 5' and 3' ends.

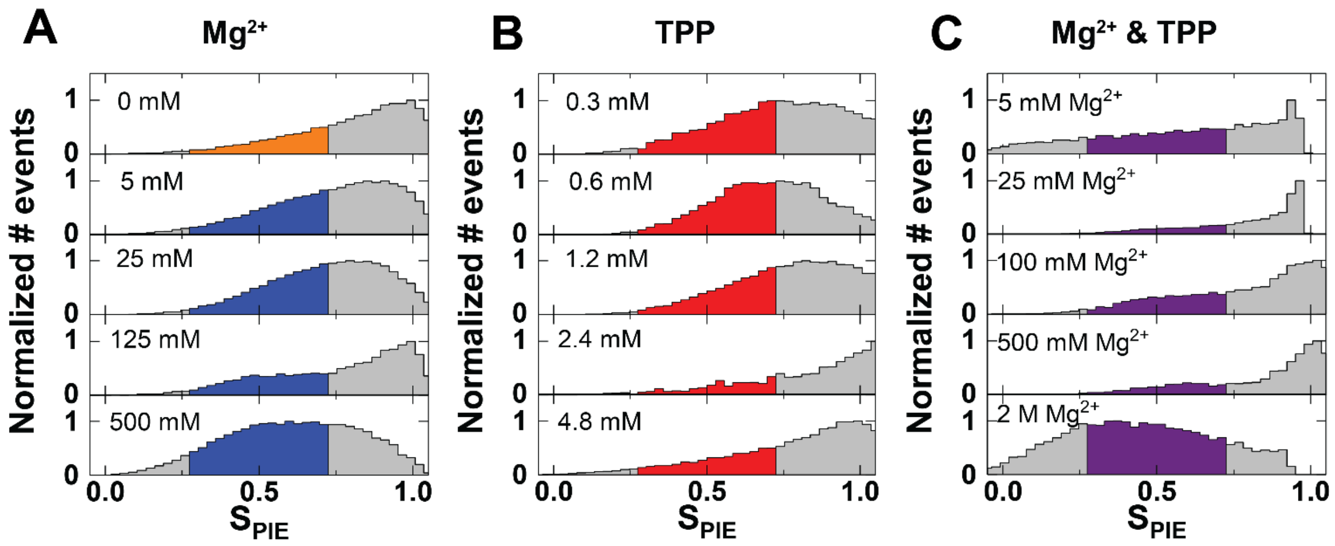


**Figure S2.** MFS Analysis was restricted to bursts with values of $S_{PIE}$ between .3 and .7 to further reduce the number of molecules analyzed with inactive acceptor and to further ensure 1:1 donor to acceptor stoichiometry. Colored regions of each plot represent the selected data which was used for analysis in Figure 3.

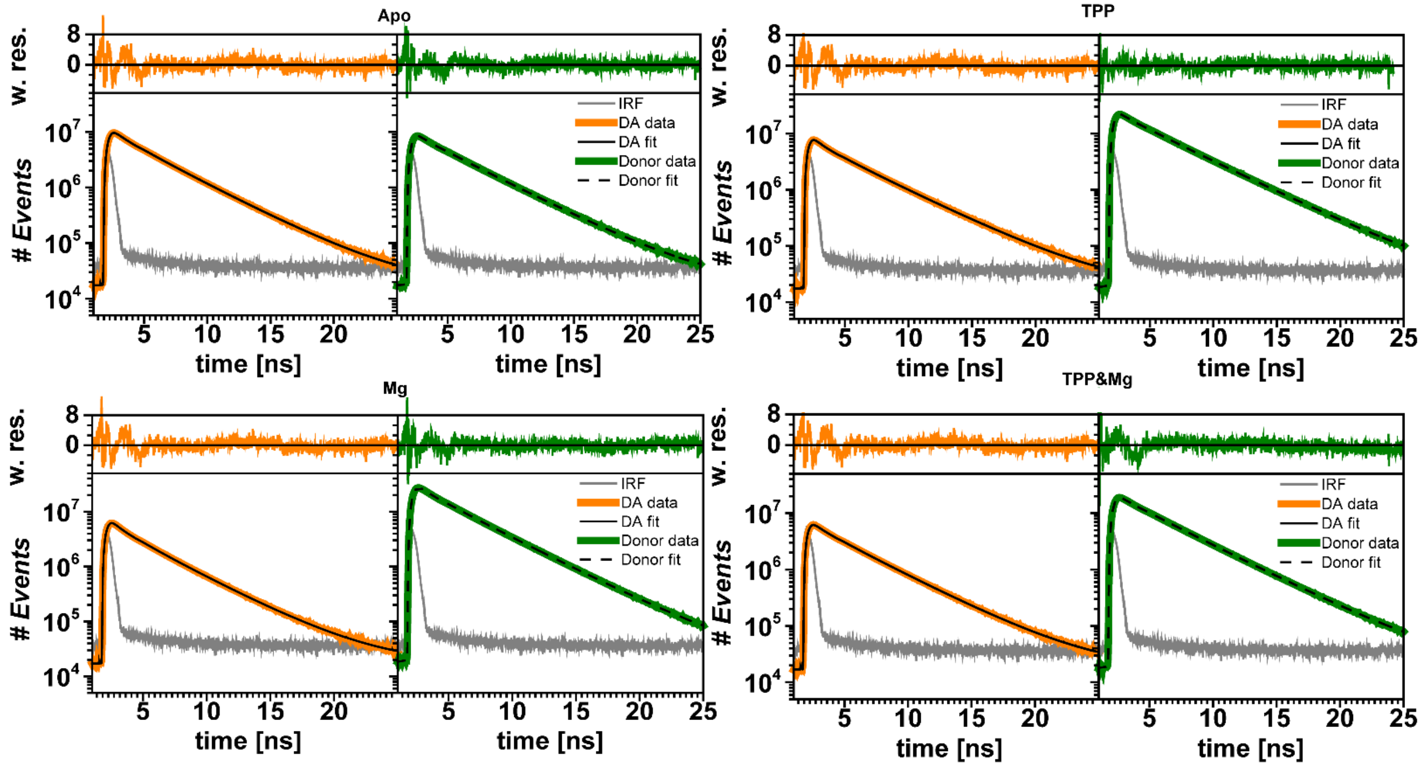


**Figure S3.** Time-resolved fluorescence decays in each experimental condition. Each fit model term corresponds to a state with Gaussian-distributed interdye distance described by the donor fluorescence lifetime of that term. Each DA curve is fit with a no-FRET decay term in addition to two FRET-exhibiting decay terms.

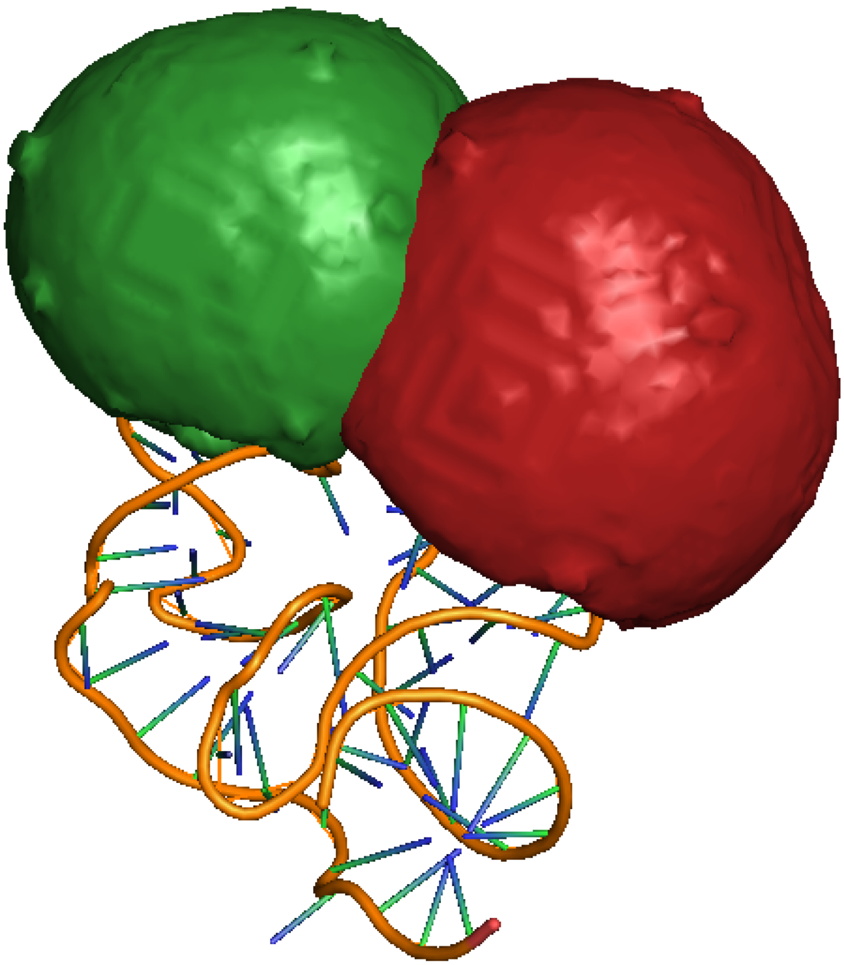

**Figure S4.** Accessible volumes from AV simulations of donor and acceptor for the TPP riboswitch closed state from the crystal structure. The donor Alexa-488 AV is shown in green, while the acceptor Cy5 AV is shown in red.

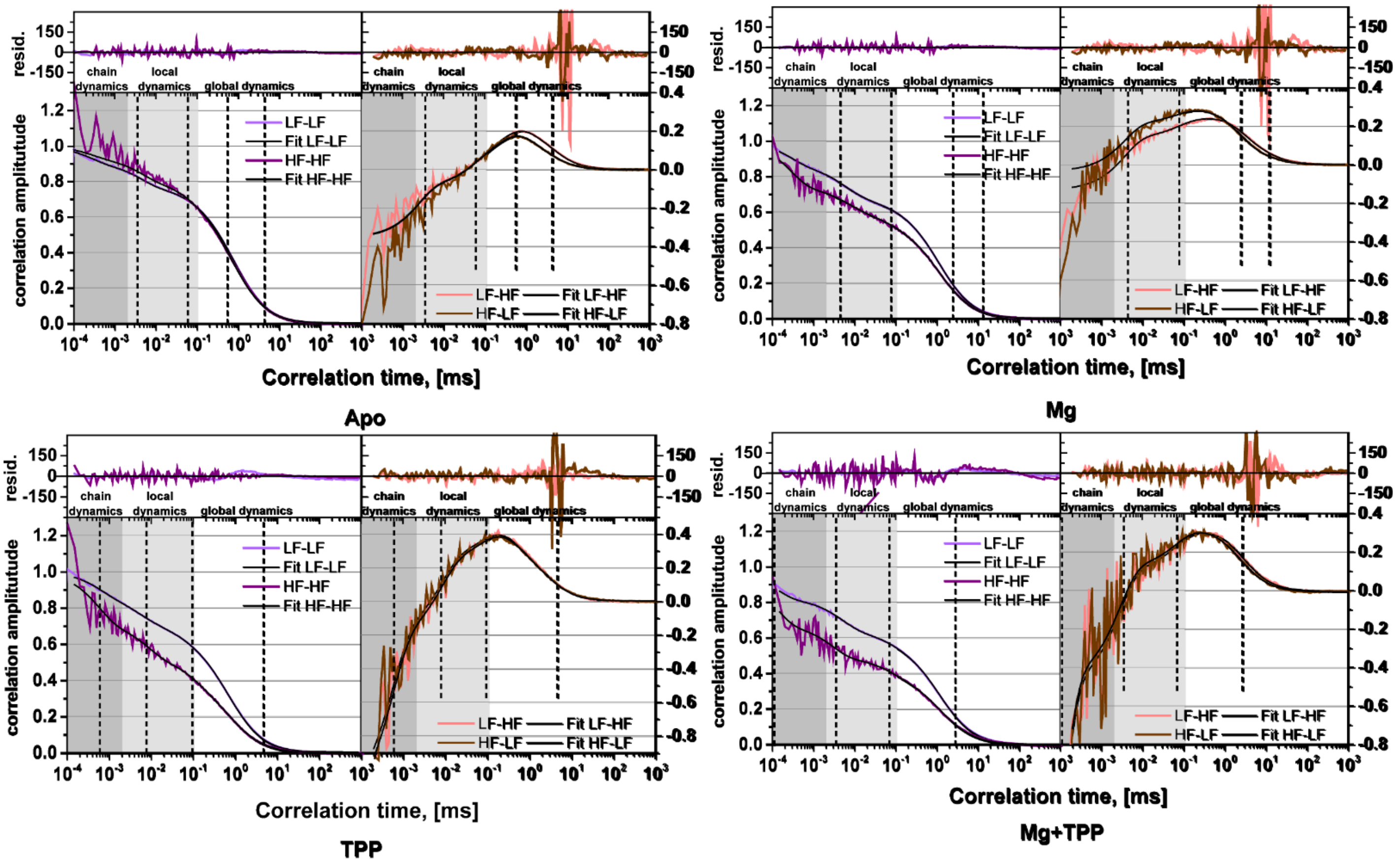


**Figure S5.** Filtered FCS Species auto and cross-correlation (sACF and sCCF) function for TPP riboswitch in a variety of experimental conditions.

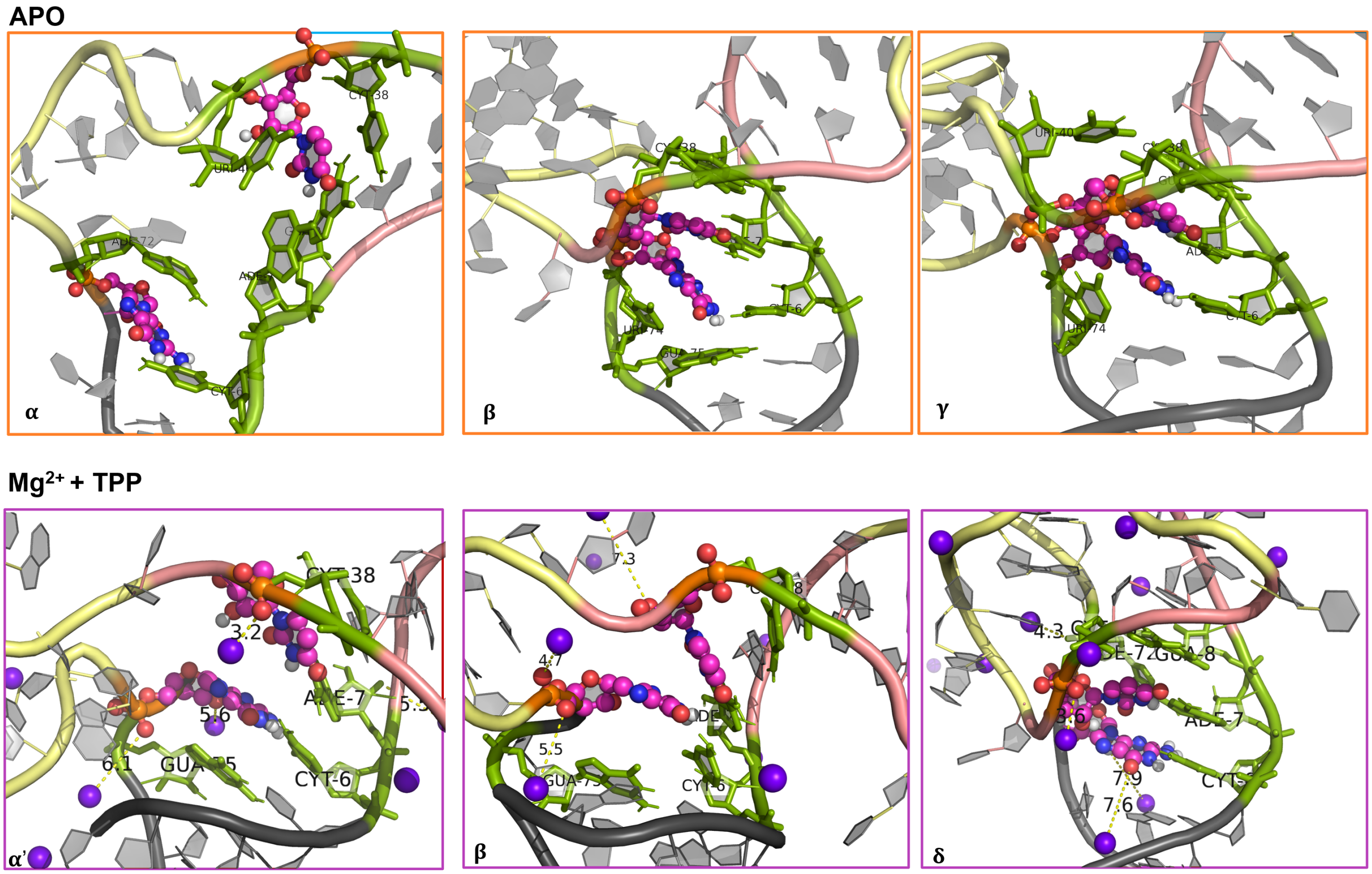


**Figure S6.** Snapshots depicting the co-stacking of U39 and G73 bases in APO (top panel, α, β, γ) and $Mg^{2+}$&TPP conditions (bottom panel, α', β, δ). The $Mg^{2+}$ are shown as purple spheres, nucleotides U39 and G73 are shown in ball-and-stick representation, and the neighboring bases are colored in green.

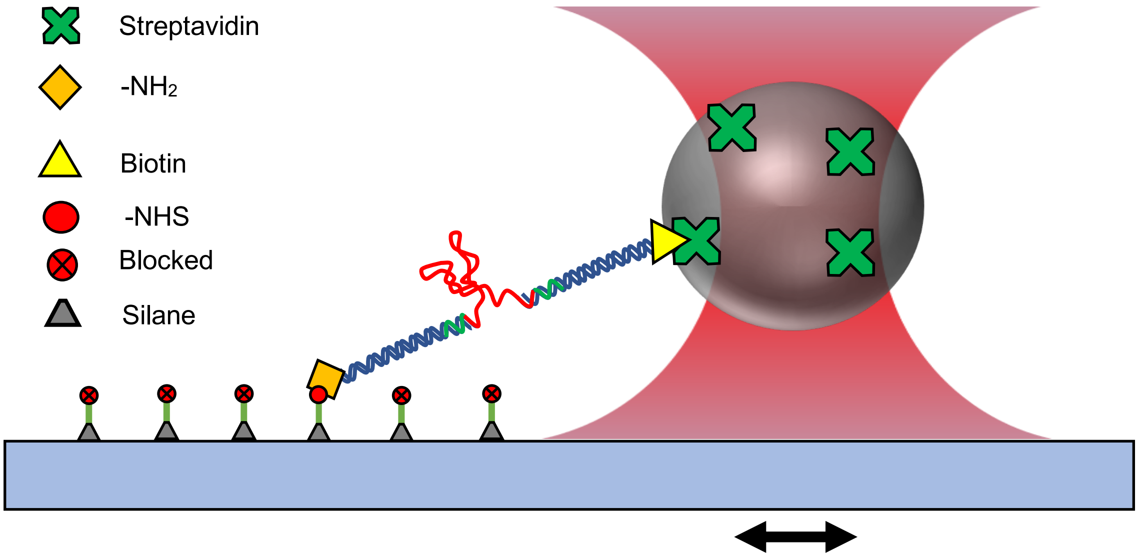


**Figure S7.** Schematic of riboswitch stretching and passive force dynamics optical tweezer experiments (not to scale). The $NH_2$-modified end of DNA-TPP riboswitch construct is anchored to the previously silanized coverglass using NHS-amine chemistry. The biotin-modified end is linked to a streptavidin-coated bead, which is trapped by the optical tweezer. The coverslip is translated relative to the fixed trap pulling the riboswitch construct. The pulling force is calculated using the calibrated trap stiffness and the displacement of the bead center from the center of the optical trap. The stretching displacement is calculated using the known stage translation, the displacement of the bead from the center of the optical trap, and the geometry of the DNA-TPP riboswitch/bead construct.

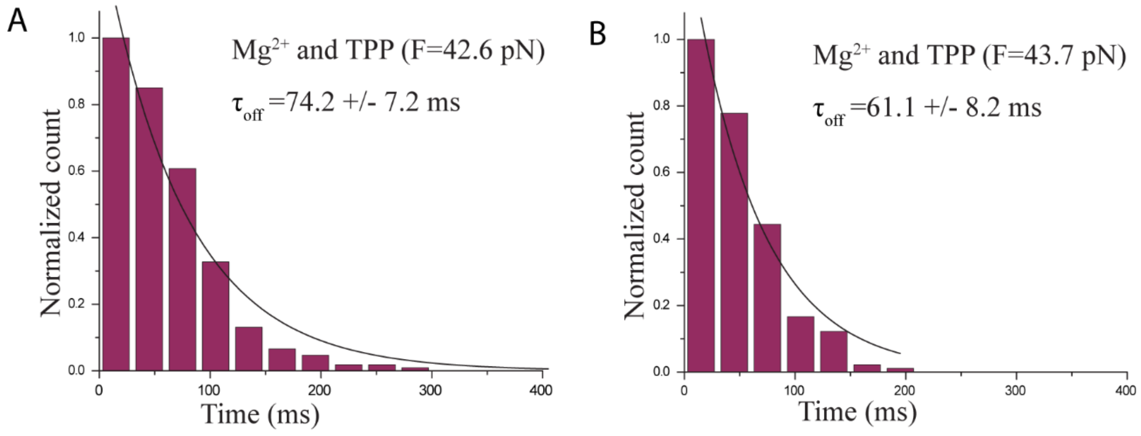


**Figure S8.** Histogram of riboswitch folded state dwell time measured in the presence of 4.8 mM TPP and 0.5 M $Mg^{2+}$ at (A) 42.6 pN, N=309, and (B) 43.7 pN, N=319. The characteristic dwell times were $\tau_{off} = 74.2 \pm 7.2$ and $\tau_{off} = 61.1 \pm 8.2$ (single exponential decay fit constant $\pm$ std. error of the fit) respectively.
